# Self-supervised generation of realistic training data enables nanoscale localization in challenging conditions

**DOI:** 10.1101/2025.07.16.665148

**Authors:** Ofri Goldenberg, Tal Daniel, Dafei Xiao, Yael Shalev Ezra, Onit Alalouf, Yoav Shechtman

**Affiliations:** Department of Biomedical Engineering, Technion–Israel Institute of Technology, Haifa, Israel; Carnegie Mellon University, Pittsburgh, PA, USA; Russell Berrie Nanotechnology Institute, Technion–Israel Institute of Technology, Haifa, Israel; Department of Electrical and Computer Engineering, Technion–Israel Institute of Technology, Haifa, Israel

**Keywords:** Simulation-to-experiment gap, Single-molecule localization microscopy, Physics-informed generative model, Self-supervised learning

## Abstract

Localization microscopy has overcome the diffraction limit, i.e. the conventional resolution limit of a microscope, enabling nanoscale biological imaging by precisely determining the positions of individual emitters such as single fluorescent molecules. However, the performance of deep learning methods, commonly applied to these tasks, depends significantly on the quality of training data, typically generated through simulation. Creating simulations that perfectly replicate experimental conditions remains challenging, resulting in a persistent simulation-to-experiment gap. To bridge this gap, we propose a physics-informed generative model leveraging self-supervised learning directly on experimental data. Our model extends the Deep Latent Particles (DLP) framework by incorporating a physical model of the Point Spread Function (PSF; the image of a single point source in the microscope) into the decoder, enabling it to disentangle learned realistic environments from emitters. Trained directly on unlabeled experimental images, our model intrinsically captures realistic background, noise patterns, and emitter characteristics. The decoder thus acts as a high-fidelity generator, producing fully labeled, realistic training images with known emitter locations. Using these generated datasets significantly improves the performance of supervised localization networks, particularly in challenging scenarios such as complex backgrounds and low signal-to-noise ratios. We demonstrate our approach on a variety of experimentally measured microscopy data, including super-resolution imaging in 2D and 3D and particle tracking in live cells, showing substantial improvements in localization precision and emitter detection.

## Introduction

Determining the precise positions of single molecules or particles in optical microscopy images is a fundamental capability enabling insights at the nanoscale, with applications ranging from biology to materials science. Single-molecule localization microscopy (SMLM) techniques, including PALM^1^, STORM^2^, and PAINT^3^, use sequential localization of emitters to reconstruct images that overcome the diffraction limit, allowing detailed biological visualization^4^. Single-Particle Tracking (SPT) utilizes localization over time to determine the trajectories of moving molecules or nanoparticles, revealing information about their dynamics, interactions, and microenvironments^5^. Extending localization microscopy to three dimensions (3D) typically requires physical modification of the microscope; a common approach involves Point Spread Function (PSF) engineering, where optical elements modify the PSF to encode depth^6–9^ or other information, such as spectral properties^10,11^ or molecular orientation^12,13^.

In localization microscopy, each emitter, e.g. a fluorescent molecule or a nanoscale particle, appears as a PSF, namely, the image of a point source formed by the microscope. In 2D SMLM the PSF is the standard, approximately Gaussian diffraction-limited spot, and localization amounts to estimating its lateral (x, y) center. Recovering the axial (z) position can be done based on PSF engineering, such as the optimally photon-efficient Tetrapod PSF^8,9^ (Figure 1a). Figure 1b shows an example application of 3D localization: fluorescently labeled DNA loci in fixed yeast cells, where each cell contains a single locus whose Tetrapod-shaped image encodes its 3D position. The goal here would be to determine the 3D positions of all loci in the field of view from a single 2D image.

**Figure 1:**
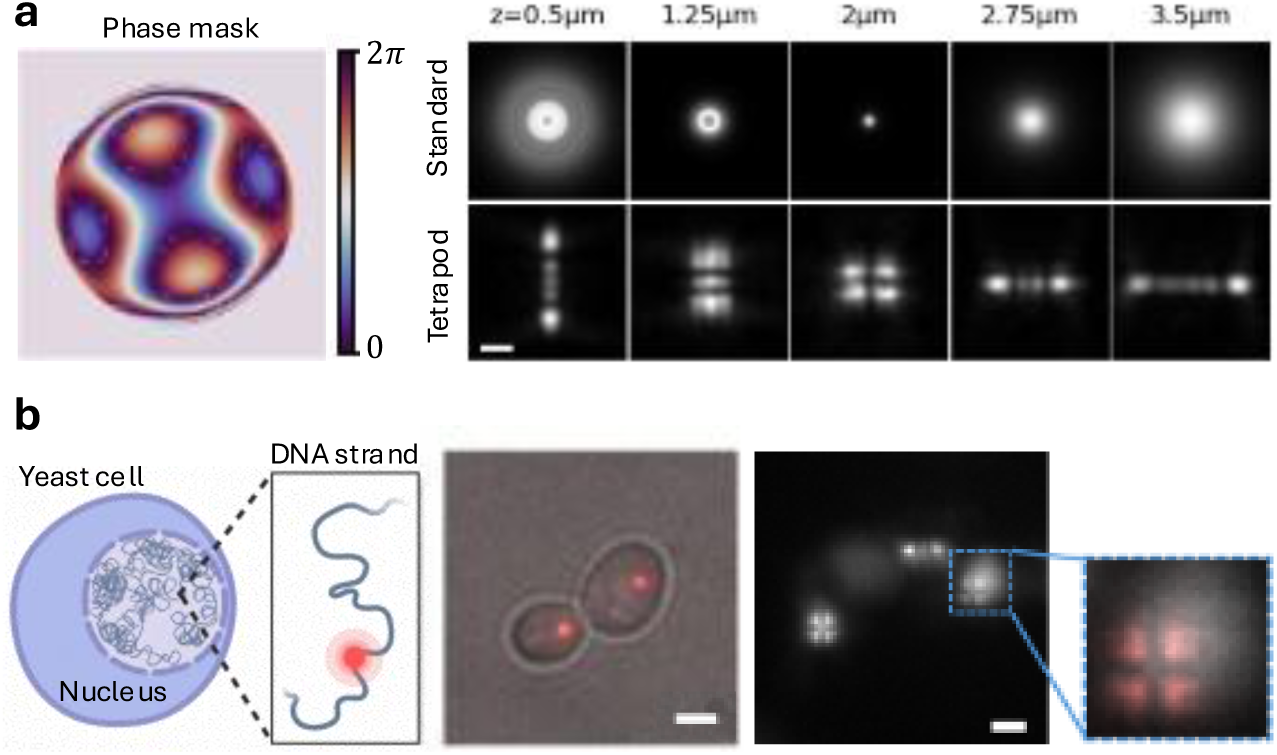
(a) Left: Model of a Tetrapod phase mask placed in the microscope’s pupil plane encodes the emitter’s axial position in the shape of the PSF. Right: measured Tetrapod PSF as a function of emitter axial position *z*, compared to the standard diffraction-limited PSF. Scale bar, 1 µm. (b) Biological sample: fluorescently labeled DNA loci in fixed yeast cells, where each cell contains a single labeled locus within the 3D volume of the nucleus (left: illustration; middle: brightfield overlay, Scale bar 1 µm). Adapted from ref^14^. The fluorescence image (right, Scale bar 2 µm) shows Tetrapod PSFs encoding the 3D positions of several emitters (inset: a single locus).

Deep learning methods have become essential tools in microscopy and SMLM, enabling powerful analysis and reconstruction^15,16^. They have been applied to various challenges, including the localization of dense emitters^17–19^, the classification of emitter color^20^, the determination of molecular orientation^21–23^, and the facilitation of end-to-end optimization of imaging systems, e.g. designing the PSF together with the reconstruction algorithm^14,18,20,24^. Importantly, the performance of deep learning approaches, particularly supervised methods, depends on the quality and quantity of the training data. Such training data is often generated via simulation, which, in the case of localization microscopy, relies on modeling the PSF and other experimental factors, because acquiring large, accurately annotated experimental datasets for 3D localization microscopy is practically impossible.

However, building simulations that perfectly mimic experimental conditions is a major challenge^25^. Accurately capturing complex, spatially varying background, realistic noise statistics, and system-specific aberrations requires significant effort and in many cases is impractical. This leads to a persistent simulation-to-experiment (sim2exp) gap, where models trained on synthetic data underperform when applied to real measurements. Furthermore, even when an adequate simulation model exists, modifying it to reflect new experimental conditions can require time-consuming manual tuning or substantial redevelopment efforts.

Recent large vision foundation models, including adaptations for microscopy^26–28^, show promise for tasks such as segmentation, cellular mapping or classification, but are ill-suited to sub-pixel localization: they are usually pre-trained on different domains and cannot be readily fine-tuned for SMLM given the scarcity of labeled data, especially across diverse SNRs and engineered PSFs. In the domain of data generation, diffusion models were adapted to synthesize realistic super-resolution images for augmentation^29^, but the generated data is typically unlabeled, and thus cannot be used for training localization networks. Other generative approaches, such as Generative Adversarial Networks (GANs)^30–33^, also attempt to bridge the sim2exp gap by learning mappings from simulated to real domains or generating images from labels. However, such post-hoc image transformations are fundamentally incompatible with the sub-pixel precision demands of SMLM. Any pixel-level modification of a simulated image can deform the PSF shape or shift its apparent center, breaking the correspondence between the image content and the original ground-truth emitter coordinates. At nanoscale resolution, where localization errors on the order of tens of nanometers are meaningful, there is no tolerance for label corruption introduced by an unconstrained generative process. Concurrently, VAE models demonstrated the ability to encode emitters’ z-position in their latent space^34^.

Here, we introduce a physics-informed VAE that embeds an image formation model directly in the generator to produce realistic, label-consistent localization-microscopy data. We propose *PILPEL – Physics-Informed Latent Particles for Emitter Localization*, an object-centric self-supervised generative model built on the Deep Latent Particles (DLP^35,36^) framework, to bridge the sim2exp gap. We extend DLP by incorporating an explicit physical PSF model into the decoder that takes encoded spatial coordinates as input. Trained directly on unlabeled experimental images, this architecture allows the model to intrinsically learn realistic background and noise patterns while disentangling the learned environment from the precise placement and properties of the emitters. The trained PILPEL model thus serves as a data generator, producing realistic, fully labeled training images with known ground truth emitter locations. This generated data is then used as highly realistic training material for supervised localization algorithms such as DeepSTORM^17^ and DeepSTORM3D^18^. We demonstrate that using these generated datasets significantly improves the performance of supervised localization networks on real-world data, particularly in challenging scenarios such as complex backgrounds and low signal-to-noise ratios. No manual tuning of simulation parameters or explicit modeling of experimental attributes is required.

## Results

Our goal is to design a controllable generative model that can generate realistic image data for the task of SMLM. In the following, we describe our proposed model that extends the DLP framework to localization microscopy.

### Physics-Informed Deep Latent Particle Model for Localization Microscopy

DLP^35,36^ is a self-supervised, object-centric image representation model, built on a variational autoencoder: an image is encoded into a set of latent “particles,” each carrying features such as 2D position and size, together with a separate background representation. These are then decoded and combined to reconstruct the input (see Methods). Training on this image-reconstruction objective, with no labels, disentangles the particles from the background. In localization microscopy, the objects of interest are individual emitters, each represented by a PSF (Figure 1). Precise localization of these PSFs is the basis for the nanoscale resolution achieved in SMLM. DLP’s explicit, position-based architecture allows fine-grained control over the objects in the generated output, making it a natural fit for our goal. We adapt DLP for localization microscopy by enabling it to recover 2D or 3D positions and intensities of point-emitters, observed as individual PSFs. We describe the model in its general 3D form; 2D localization follows as the special case in which the axial coordinate is fixed (z = 0) and the engineered phase mask is replaced by a flat-phase pupil, reducing the PSF to the standard diffraction-limited spot.

We extend the original DLP model to explicitly encode the key emitter properties, i.e., spatial coordinates ({*x*, *y*, *z*}) and intensity (*I*), such that each latent particle corresponds to a single emitter. Crucially, the model learns additional image components such as local and global background, SNR, and effective emitter density. We term this extended framework **PILPEL**: **P**hysics-**I**nformed **L**atent **P**articles for **E**mitter **L**ocalization.

This multi-component learning is achieved by incorporating a physical model of the PSF into the decoder, where the encoded {*x*, *y*, *z*} coordinates are used as input to simulate PSFs as part of the image formation process. The PSF model serves as a strong inductive prior, embedding physical constraints into the generative process. The resulting physics-informed architecture allows the model to disentangle the PSFs from their surrounding scene, including noise and measurement artifacts, while explicitly capturing the key parameters of interest, namely, emitter location and intensity, enabling fine-grained control and accurate reconstruction. The framework is agnostic to the specific PSF: adapting it to a different engineered PSF (e.g. astigmatism^6^, double-helix^7^) is straightforward and requires no architectural changes.

Our model architecture is illustrated in Figure 2: an object-centric encoder–decoder in which a differentiable physical PSF model, embedded in the decoder, drives image formation from the latent emitter coordinates. The reconstructed image is composed of separable components, namely, emitter PSFs, their local background, and a global background.

**Figure 2:**
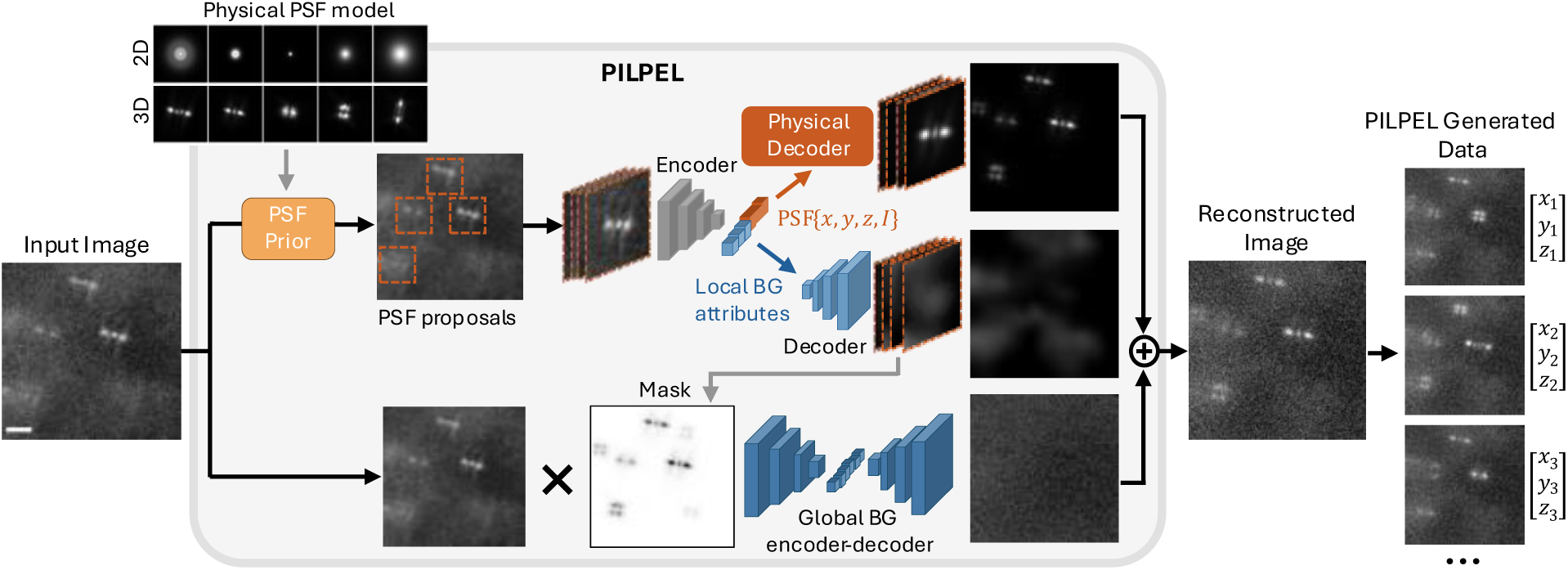
PILPEL – Physics-Informed Latent Particles for Emitter Localization. Encoding-decoding scheme with a latent representation of PSF-objects. The model divides the input image into overlapping patches and performs template matching using a precomputed PSF dictionary (standard 2D or engineered 3D PSFs, top left) to propose candidate emitter locations. Patches around these locations are then processed by an encoder that generates disentangled latent vectors encoding spatial coordinates, intensity, and appearance features of the local background. A decoder reconstructs the image using a physics-based PSF generator and a CNN-based local-background generator. These generated patches, along with a separately modeled global background, are combined to form the final reconstructed image. The entire system is trained end-to-end within a VAE framework, balancing reconstruction accuracy and latent regularization. Because the coordinates passed to the physical decoder are the ground truth of the generated image by construction, re-sampling them together with the latent attributes and passing them through the decoder provides new labeled images, with known {x, y, z} per emitter, that serve as training data for supervised localization networks. Scale bar, 2 µm.

Once trained on real measurements, the decoder of our PILPEL model serves as a high-fidelity generator of extremely realistic images with precisely known emitter positions (Figure 2, right). Importantly, because PILPEL is trained for image reconstruction rather than localization, its encoded coordinates are not precise enough to serve for super-resolved localization directly. Instead, we sample new, exactly known coordinates into the decoder’s physical layer, which are the ground truth for the generated image by construction (see Methods for the full details). These generated, labeled images inherit the realistic background and noise characteristics learned from the experimental data, as well as emitter intensities, densities, and their spatial distribution. This enables large and accurately labeled training datasets for supervised localization models, specifically DeepSTORM^17^ and DeepSTORM3D^18^ in this work. This data closely mimics real measurements, thereby narrowing the sim2exp gap while automatically capturing unmodeled experimental nuances.

In the following sections we show that the generation of realistic data improves the downstream performance of localization tasks. We first establish the method’s accuracy on controlled simulations with known ground truth, then demonstrate its impact on real biological data, consisting of one dataset of 2D localization and three datasets of 3D localization. Full experimental details are provided in the Supplementary.

### Bridging the sim2exp Gap – Controlled Simulations

To quantify performance and demonstrate the benefit of using training data generated using PILPEL, we evaluated DS3D for 3D localization across different training datasets. First, we designed a test set resembling yeast cell measurements with point sources shaped by a Tetrapod phase mask^37,38^. To simulate the typical autofluorescence observed in these cells, we added random asymmetrical 2D Gaussians around each PSF. Crucially, to reflect experimental conditions containing unknown noise profiles, we introduced complex patterns that are absent from the original DS3D simulator and are difficult to tune manually. Specifically, we simulated inhomogeneous backgrounds using Perlin noise, which is commonly employed in microscopy image analysis to model non-uniform illumination and other artifacts^19^. We term this test set PerlinData.

We then trained DS3D on three different datasets: (1) the original simulator with a wide range of SNR and background values; (2) the original simulator with an added local background (the same asymmetrical Gaussians used in the test set, practically introducing prior knowledge manually); and (3) PILPEL-generated dataset trained on the PerlinData images.

Figure 3a illustrates these datasets, showing that, as expected, the generated data most closely resembles the test set. PILPEL successfully learned Perlin noise with various levels and complex background patterns. These elements are not modeled in the original DS3D simulator and contribute to the sim2exp gap that can significantly degrade localization network performance. Consequently, DS3D trained on this generated data achieves superior performance, both in localization accuracy and in the number of detected emitters.

**Figure 3:**
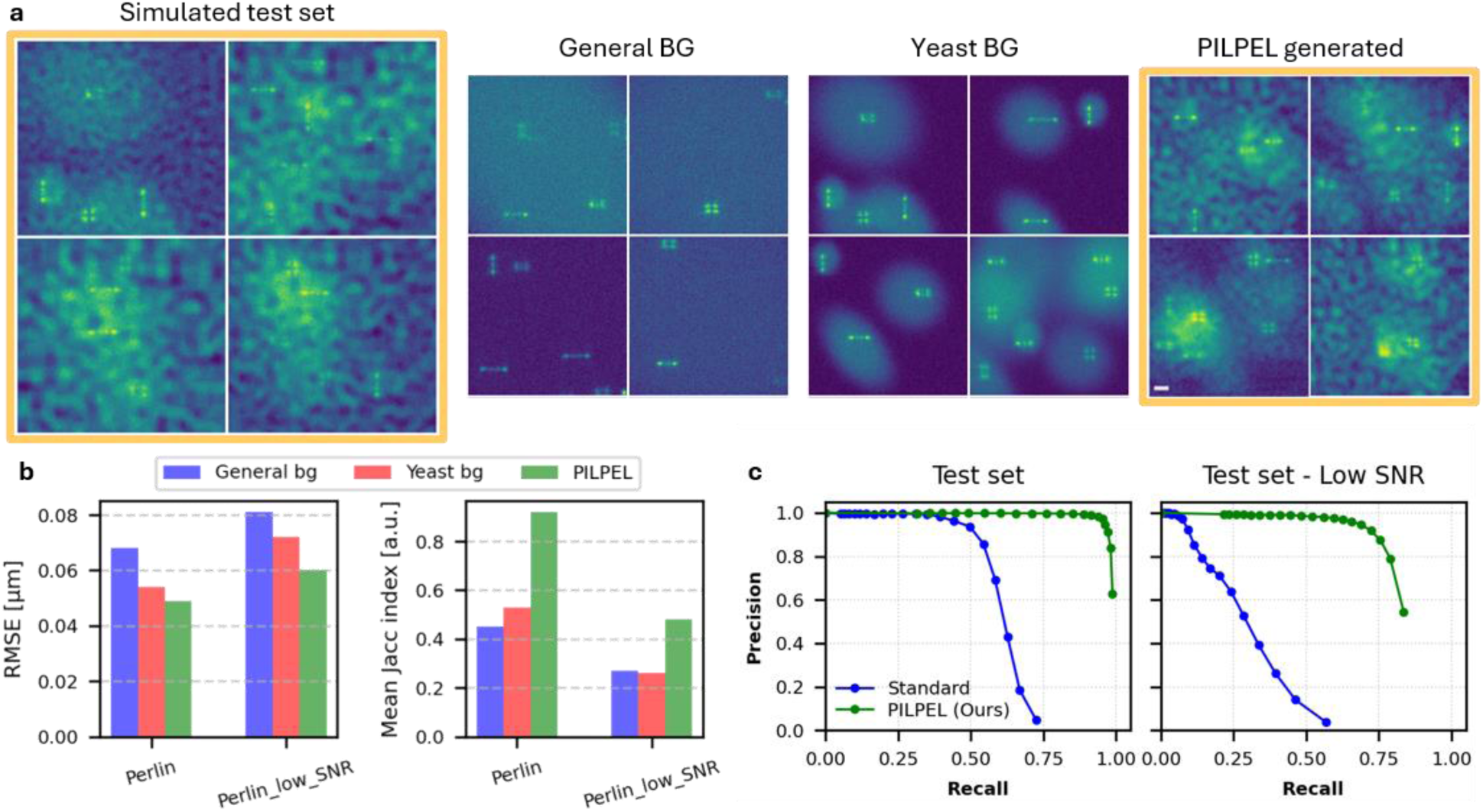
Robustness simulation – DS3D localization performance. (a) Comparison of synthetic training datasets used for DS3D localization of the Perlin test set (left). General BG – the standard background in DS3D; Yeast BG – includes local background around emitters to mimic yeast cells; and PILPEL – generated via our model. Scale bar, 2 µm. (b) Improvement of training on generated data. Localization RMSE (lower is better) and Jaccard index (higher is better) for DS3D trained on the generated dataset vs. training on yeast-BG and original-DS3D datasets. (c) Precision-Recall analysis on the PerlinData test set. PILPEL (green) demonstrates superior detection robustness compared to the standard DeepSTORM3D baseline (blue). While the baseline’s precision degrades rapidly, particularly in the Low-SNR regime, PILPEL maintains high precision (> 0.9) across a significantly wider recall range.

Figure 3b summarizes the performance gains of using the generated dataset under challenging conditions, split into PerlinData and low-SNR-PerlinData to show the benefit is even greater in low-SNR cases. Localization RMSE improves by 19 nm (a 28% improvement), and the Jaccard index (measuring the overlap between detected and ground truth localizations) doubles from 0.45 to 0.92, compared to the baseline. While introducing prior knowledge into synthetic training data improves performance (yeast background, red bar), training on generated data (PILPEL, green bar) is still significantly better, and also eliminates the need for incorporating known or assumed prior information in the DS3D data simulator.

To further evaluate the detection capability of the proposed model and its robustness against false positives, we performed a Precision-Recall (P-R) analysis on the PerlinData test set. Ground truth emitters were matched to localized coordinates using a strict lateral distance threshold of 100 nm, the detection confidence threshold varied from 0 to 800 (the range of output possibilities) to generate the curves. As shown in Figure 3c, there is a decisive performance gap between our method and the standard baseline. The baseline fails significantly on unmodeled noise, with precision collapsing after approximately 50% recall. In contrast, PILPEL maintains near-perfect precision (∼1.0) up to ∼95% recall. Notably, at a fixed high precision of 0.9, PILPEL yields approximately 2 × higher recall than the baseline. This advantage is even more pronounced in the challenging Low-SNR regime. Here, PILPEL maintains > 0.9 precision up to 75% recall, whereas the baseline fails (dropping below 0.9 precision) at only 15% recall. These results demonstrate the robustness of our approach to unmodeled noise, showing how PILPEL-generated data mitigates the sim2exp gap. See Supplementary Sec. B for extended details.

### 2D Super-resolution Reconstruction of Microtubules

Having validated our approach under controlled conditions with known ground truth, we next evaluate its performance on real-world biological data, starting with 2D super-resolution imaging of microtubules, using DeepSTORM^17^.

We used a dataset of microtubules in fixed cells, imaged via conventional 2D STORM^39^. We trained our PILPEL model directly on the experimental 2D STORM image frames. Since 2D localization requires only lateral coordinates, we fixed the axial coordinate in the PSF model to zero and used a flat pupil (zero phase mask), thus generating standard diffraction-limited 2D PSFs rather than the engineered 3D PSFs used in the other experiments. The model architecture remained unchanged, learning to represent each frame as a set of 2D point emitters with associated intensities, along with local and global background components. Figure 4a illustrates the model’s learned decomposition of measured frames into their constituent components, demonstrating its ability to accurately represent the input data. The generated labeled dataset from PILPEL was then used to train DeepSTORM. We compared three reconstruction approaches on the same dataset: (1) ThunderSTORM^40^, a classical localization algorithm based on sub-pixel fitting; (2) DeepSTORM trained using the original simulator provided with the published method; and (3) DeepSTORM trained on our PILPEL-generated dataset.

**Figure 4:**
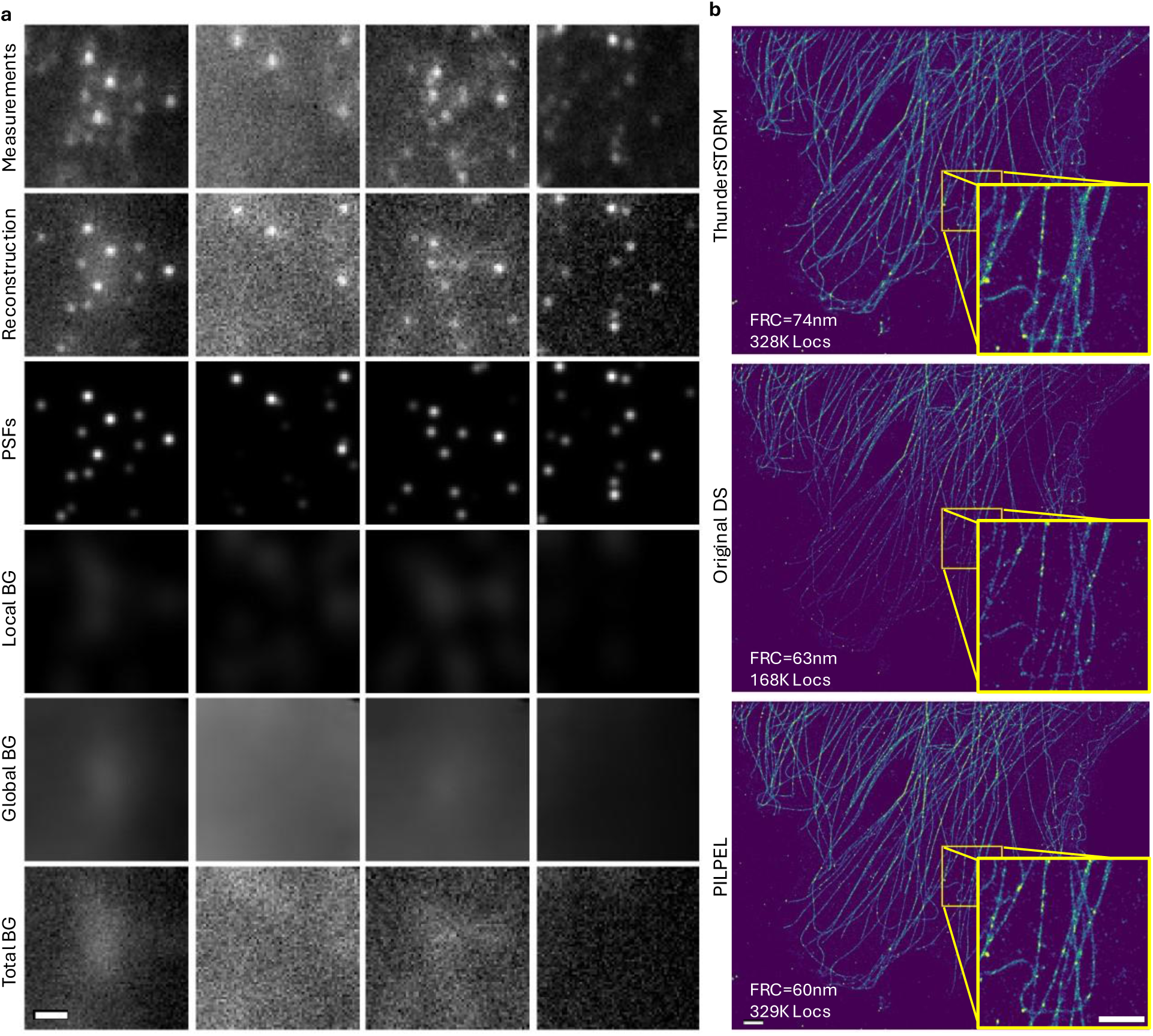
(a) 2D STORM experimental frames and PILPEL output components. The first row shows five measured images, the second row shows PILPEL’s final reconstructions. *The remaining rows show PILPEL’s reconstruction components:* the PSF component (third row) representing the isolated emitter signals; the local background component (fourth row) around each emitter; the global background component (fifth row), representing the general background across the entire field of view; and the total background which is the sum of all non-PSF components, including the added noise. *Scale bar, 2 µm.* (b) 2D super-resolution reconstruction comparison of ThunderSTORM, original DS, and DS trained on PILPEL-generated data. Compared to ThunderSTORM, original DS better recovers dense structures, and PILPEL-trained DS further enhances the reconstruction. Scale bars, 3 µm (main), 2 µm (inset).

Figure 4b presents the super-resolution reconstructions of a microtubule structure. The original DeepSTORM (trained on simulated data) achieves sharper reconstructions compared to ThunderSTORM, especially in dense regions with overlapping PSFs. However, it still underperforms in certain challenging areas (e.g., lower left region), typically corresponding to low SNR regions or areas with characteristics that differ substantially from the training data. DeepSTORM trained on PILPEL-generated data demonstrates significantly improved performance in these problematic areas while further enhancing the overall reconstruction, maintaining sharp, resolved details (see Figure 4b insets) and improving visibility, thus outperforming both ThunderSTORM and the original DeepSTORM trained on simulated data.

The total number of localizations increased from ≈168K for the original DeepSTORM to ≈329K for the PILPEL-trained version across 600 frames (530×358 pixels), representing a 1.96-fold increase. This improvement translates to both enhanced spatial resolution and more complete structural representation of the microtubule network. To empirically validate these results, we used Fourier Ring Correlation (FRC), filament structural analysis, and spatial distribution analysis, as detailed in Supplementary Sec. C.

### 3D Super-resolution Imaging of Mitochondria

Having demonstrated PILPEL in the 2D setting, we now turn to the more challenging 3D case, where depth is encoded through an engineered PSF. We apply the method to a 3D super-resolution dataset of mitochondria^41^ imaged with the Tetrapod PSF. As shown in Figure 5a, training on our generated data dramatically improves the final reconstruction. The number of detected localizations increased from 119K to 581K (∼ 4.9 × increase) across 20K frames. This translates to improved image quality, i.e. fuller features (as seen in Figure 5b), and potentially faster acquisition. Figure 5c shows measured frames and corresponding PILPEL reconstructions.

**Figure 5:**
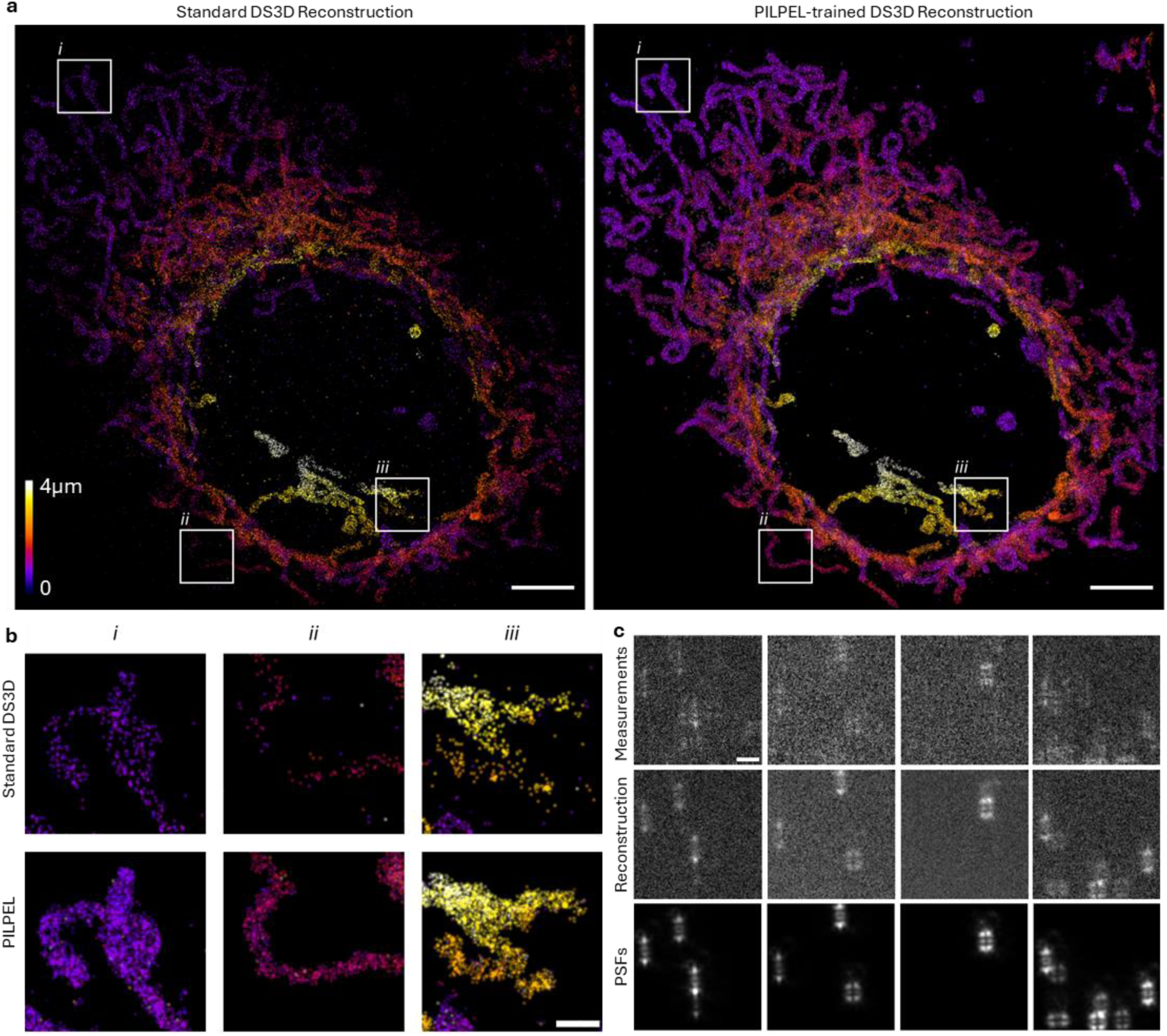
3D super-resolution reconstruction of experimental data. (a) Super-resolved reconstructions of mitochondria (thickness ≈ 4 µm), rendered as 2D histograms with color-coded *z*-positions, using DS3D trained on the original simulated data (left) versus our generated data (right). Scale bar, 5 µm. (b) Magnified views of regions i, ii, and iii as indicated in (a). Scale bar, 1 µm. (c) Examples of randomly sampled measured images, accompanied with their matching PILPEL reconstruction generated by our model, used to train DS3D. Bottom row shows the PSFs component of the reconstructions, defined by {*x*, *y*, *z*} coordinates, that serves as the GT training labels. Scale bar, 3 µm.

A possible concern would be that an increase in detections may be associated with a higher rate of false positives. We evaluate this with a volumetric density analysis on 20 background ROIs compared to 20 volume-matched on-structure ROIs (Supplementary Sec. C). While PILPEL’s off-structure density is indeed higher than DS3D (4.56 vs. 0.45 loc/µm³) this absolute increase (4.11 loc/µm³) is negligible compared to the signal gain on the biological structure (635.5 loc/µm³). Furthermore, these residual localizations can be suppressed by applying a stricter confidence threshold, made possible by the higher total number of detections, or by using standard density filters commonly applied in SMLM reconstruction^40^. For the visualization in Figure 5, the reconstruction from our method was post-processed with such a filter to match the original’s noise level, while the original reconstruction was left as-is, as filtering would degrade the sparse biological structure. Crucially, the high detection rate achieved by our method makes the application of such filters feasible. While our dense reconstructions survive filtering and preserve the underlying structure, the baseline’s sparse detections vanish under the same processing. The unfiltered version is available in Supplementary Figure 8.

Importantly, our method learns directly from the measurements without the need for manual tuning of the DS3D simulation parameters, which requires hours if not days, depending on user expertise. Human capability to verify and check vast amounts of images is limited, whereas PILPEL learns to generate based on the entire dataset, providing more comprehensive optimization. Furthermore, manual tuning relies on subjective visual assessment (“does it look similar enough?”), while PILPEL employs computationally optimized image similarity loss between measurements and generated data, which is more efficient and robust. While recent work aims to automate this process^42^, such approaches are still limited by the simulator’s explicit model and pre-defined parameters, which do not capture unmodeled traits that our generative approach learns directly.

### 3D Localization of DNA Loci in Yeast Cells

Beyond super-resolution microscopy, we further assess the generalizability of our approach in a distinct biological system characterized by local background features not modeled in the original DS3D simulator. We apply the same pipeline to experimental images of fluorescently labeled DNA loci in fixed yeast cells^38^, where each cell exhibits a single fluorescent spot shaped by the Tetrapod phase mask. In such a biological system, an example application would be tracking loci over time in 3D under different conditions^5,38,43^; here, we use it to demonstrate our method under a noise regime distinct from STORM.

Dense low-SNR measurements were used as a challenging dataset for DS3D localization, comparing a model trained with the original simulator versus a model trained on generated data derived via our data generation pipeline. Figure 6a shows the measured dataset and the corresponding PILPEL reconstructions. Figure 6b displays the measured images alongside localization results from both localization networks. The general detection rate increased by ∼ 1.47 ×, from 12,561 detections with the original net to 18,461 detections using the PILPEL-generated dataset, across three time-lapse measurements (100 time points each) in a 1200 × 1200 FOV. In addition to that, mean detection confidence values increased by 40% using PILPEL.

**Figure 6:**
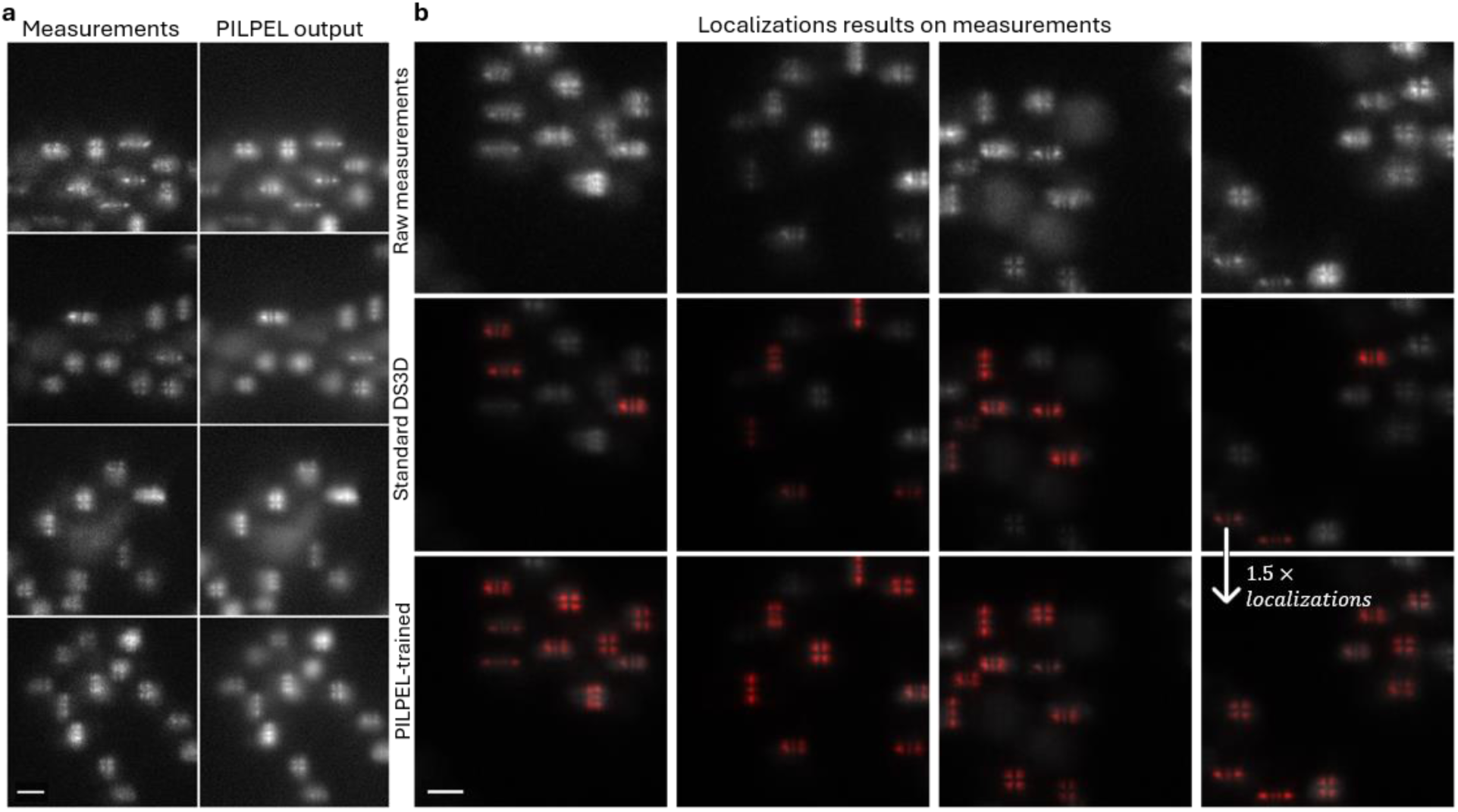
Fluorescently labeled DNA loci in yeast cells. (a) PILPEL trained on experimental data, showing dense yeast cells with pronounced, spatially varying local background. Left column is the input – raw measured images; on the right is the output – the reconstructions generated by the model. Reconstructions capture cell-specific background and noise patterns, illustrating that the model learns biological structures directly from the data. (b) Comparison of localization results. Localization overlays on the same measurements (white) with 3D detections rendered (red). Middle row: DS3D trained on standard simulated data; bottom row: DS3D trained on PILPEL-generated data. Evidently, the PILPEL-trained network yields visibly higher red-on-white overlap and recovers more emitters missed by the standard baseline. Scale bars, 3 µm.

Ground-truth positions are unavailable in such experiments, but standard GT estimation in single cells is MLE localization – fitting xyz coordinates of the PSF model together with asymmetric Gaussian background. This is relevant when the model matches the measurement, and MLE is not always correct in such difficult cases. Hence, we conducted a small experimental test set of 1,500 single-cell measurements with relatively high SNR where we assume the MLE fitting will work, and compared the results to the two networks’ localizations. Localization RMSE improves from 107 nm to 96 nm, a 10% improvement, using PILPEL. Importantly – this test set comprised high SNR data, whereas a more significant improvement will be on low SNR cases, where the model deviates substantially from the measurement. The advantage of our method is that it generalizes the model and can identify more emitters, in a large FOV with multiple emitters. This is reflected in greater detection rate, seen also on the provided images, including low-SNR emitters missed by the original DS3D.

Compared to the synthetic yeast-BG dataset from Figure 3, our model’s ability to learn unmodeled features is clearly demonstrated. While one can incorporate local background to the DS3D simulator, it still does not adequately resemble the actual measurements seen in Figure 6. PILPEL’s generated data improves this correspondence and removes the need for manual implementation, allowing its use for future datasets with different noise patterns and shapes.

## Discussion

High fidelity data simulation is crucial for all simulation-based learning. In this work we pushed data simulation in localization microscopy to the extreme realistic limit. This is achieved by integrating an explicit PSF model within a self-supervised, object-centric framework that disentangles emitters from complex background and noise. As our benchmarks demonstrate, training supervised models on this generated data significantly enhances localization accuracy and detection rates, especially for challenging or novel experimental conditions. Furthermore, this approach eliminates the need for manual simulation tuning and the explicit modeling of complex image elements, offering a robust, adaptable solution for improving deep learning performance in microscopy.

To further support the generalizability of our method, we applied PILPEL to an additional 3D dataset of bacteria using a different depth-encoding PSF (astigmatism). The results (Supplementary Sec. A) demonstrate the applicability of PILPEL to another distinct biological sample, background profile, and SMLM modality.

Despite the advances demonstrated, our method exhibits several limitations. Training the generative model requires additional computational time compared to training on synthetic datasets, although it significantly reduces manual human intervention (see Supplementary Sec. F for computational details). Additionally, accurate reconstruction still necessitates a PSF model. This is obtainable either by a numerical model or, better, from experimental calibration (see Supplementary Sec. D for robustness analysis).

Future work includes leveraging this generative capability for a variety of microscopy scenarios that are hard to simulate, e.g., particle flows, or living organisms, extending to dynamic tracking via DDLP for video^36^, and exploring latent space manipulation for tailored dataset generation for downstream tasks (e.g., denoising). Another promising direction is to incorporate learned aberrations^44^, enabling also the optimization of the PSF model itself.

Overall, PILPEL replaces hand-crafted simulation with a generative model that learns the experiment itself, producing realistic, labeled training data automatically, extending the applicability of localization microscopy to increasingly challenging experimental datasets.

## Methods

### Downstream localization networks

In this work we demonstrate the use of PILPEL-generated data to train supervised localization networks, DeepSTORM for 2D and DeepSTORM3D for 3D localization, both of which conventionally rely on simulated training data. DeepSTORM (DS^17^) is a deep convolutional neural network (CNN) method designed for high-density 2D SMLM that reconstructs super-resolved images from diffraction-limited frames of blinking emitters. DeepSTORM3D (DS3D^18^) is a CNN-based method for dense 3D SMLM over large axial ranges. It analyzes 2D images containing overlapping engineered PSFs (such as the depth-encoding Tetrapod PSF, Figure 1a) and outputs a 3D localization map. These supervised networks are trained entirely on simulated data, where large datasets are generated using a physics-based model of the microscope and PSF. These simulations incorporate realistic experimental variability, including different emitter densities, signal-to-noise ratios, background levels, and noise models. A significant step in the analysis pipeline is the generation of training data that resembles the experiment. This involves significant manual tuning and can be highly time-consuming even for an experienced user. Our model, among other contributions, alleviates this manual tuning completely.

### Deep Latent Particles (DLP)

Deep Latent Particles (DLP^35,36^)^*^ is a self-supervised, object-centric image representation model that decomposes an input image 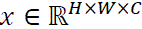 into a set of *K* structured foreground latent particles 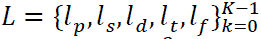. Each particle encodes the following attributes: *l_p_* ∈ ℝ^2^ denotes the 2D position, *l_s_*∈ ℝ^2^ the spatial scale (bounding box size), *l_d_*∈ ℝ the appearance order, *l_t_*∈ ℝ the transparency, and *l_f_*∈ ℝ*^m^* the latent visual features. A key component of DLP is the spatial prior, where particle positions are determined using a spatial softmax (SSM) operator applied to activation maps from a patch-based convolutional network. In addition to the *K* foreground particles, the model includes a single background particle *l*_bg_ ∈ ℝ*^m^*^bg^ to represent the background visual features. The values of *K*, *m*, and *m*_bg_ are model hyperparameters. To learn these attributes, the model decodes the latent particles into glimpses and composes them together with the background to reconstruct the input image. Training is performed end-to-end using a variational autoencoder (VAE^45^) framework, with the objective of maximizing the evidence lower bound (ELBO), combining image reconstruction loss with a KL-divergence regularization term. The disentangled latent representations learned by DLP have been successfully applied to a variety of downstream tasks, including video prediction^36^, reinforcement learning^46^, and imitation learning for robotics^47^. For a detailed overview of the model, we refer the reader to the original work^36^.

### PILPEL architecture and training

In our modified model, the physics-informed prior replaces the spatial softmax (SSM)-based prior used in the original DLP to propose keypoint locations. Specifically, we employ a physical model of the 3D PSF that accounts for the optical parameters used in the experiment. This PSF model can be theoretically derived or experimentally calibrated, for example, by using phase retrieval methods such as VIPR^48^. Similar to DS3D^18^, our model includes a differentiable optical layer that simulates image formation from emitter coordinates to the final sensor image. The same PSF model is used both in the prior module to propose emitter locations and in the decoder to generate the PSFs of each reconstructed patch.

As illustrated in Figure 2, the input image is initially divided into overlapping patches. Each patch undergoes a 3D template matching process against a dictionary of pre-calculated PSFs spanning the expected axial *z* range (for 2D localization, the dictionary reduces to the single in-focus PSF and matching is purely lateral). This yields an initial estimation of potential PSF positions (*l_x_*, *l_y_*, *l_z_* coordinates) within each patch. The maximum cross-correlation score is used as a confidence metric to filter out patches unlikely to contain any PSFs, resulting in a proposed set of latent particles.

Following this coarse-grained proposal stage, we extract image patches centered around each proposed PSF location. These patches, intended to capture both the PSF and its local surroundings, serve as input to a particle encoder module. The encoder outputs the latent attributes for each proposed object. These are disentangled by construction, as detailed below, with specific latent attributes representing subpixel offsets (*l_dx_*, *l_dy_*, *l_dz_*) to refine the initial position estimate, the PSF’s intensity *l_I_*, and additional latent attributes encoding the appearance features of the patch. The intensity factor *l_I_* allows the model to capture experiment-specific SNR and brightness distributions and enables an automatic pruning mechanism (by driving the factor to zero during optimization), which adapts to varying emitter densities by removing unnecessary proposals.

The decoder reconstructs the image from these latent representations and consists of two distinct components. The first is the differentiable physical layer that takes the refined spatial coordinates (initial estimate and the learned offset, *l_x_*+ *l_dx_*, etc.) and the intensity factor from the latent vector *l_I_* to generate a PSF image. The second component is a CNN-based decoder which takes the remaining latent appearance attributes and generates a corresponding image patch representing the local background surrounding the PSF. These are combined and placed into the final reconstructed image at their corresponding global coordinates.

To learn global background characteristics, the generated object patches are used to mask the input image. This masked input is processed by a separate encoder-decoder network dedicated to modeling and reconstructing the general background component of the scene. An additional encoder learns a latent attribute *l_n_*, used as the standard deviation of random noise added to the general background.

Finally, the generated components (PSFs, local background patches and general background) are added together to produce the final reconstructed image. The entire model is optimized end-to-end within the VAE framework by minimizing the ELBO – the reconstruction loss (typically MSE or a perceptual loss) between the output image and the original input image, and KL-divergence regularization term for the stochastic latent variables. An ablation study evaluating the contribution of each module is provided in Supplementary Sec. E.

### Generating labeled training data

To produce labeled training data, we encode an input image, sample new PSF coordinates (drawn randomly around the original encoded coordinates), and pass them through the decoder to generate realistic perturbations of the input image (Supplementary Figure 9). The model reproduces the global and local background characteristics learned from the real image, while the newly assigned PSF coordinates serve as the training labels. These synthetic yet realistic images, paired with their exact emitter positions, are then used to train a supervised localization network.

Notably, since PILPEL’s objective is image similarity rather than localization accuracy, the encoded coordinates are not guaranteed to reach the sub-pixel precision required for single-molecule localization: the reconstruction is visually faithful to the input, but the encoded PSF positions may be slightly shifted from the true ones, with a discrepancy of approximately 90 nm 3D-distance RMSE in simulation – too large to use directly as ground-truth labels. However, this potential inaccuracy is irrelevant for our purpose, since the decoder is driven by new, precisely known coordinates fed into its physical layer, which by definition constitute the ground truth for the newly generated image.

### Experimental datasets, sample preparation, and imaging

#### 2D STORM (microtubules)

This dataset, originally published in Saguy et al.^39^ consist of fixed COS7 cells immunolabeled for α– and β-tubulin (Alexa Fluor 647), imaged by STORM with an oil-immersion objective (NA = 1.49, 100× magnification), as described in details in ref.^39^.

#### 3D STORM (mitochondria)

This dataset, originally presented in Orange-Kedem et al.^41^, consists of 3D STORM imaging of mitochondria in fixed COS7 cells labeled with Alexa Fluor 647. Imaging used a 4f Fourier-plane extension equipped with a Tetrapod phase mask encoding an axial range of 4 µm, and an oil-immersion objective (NA = 1.35, 100× magnification). The experimental PSF was retrieved with VIPR^48^ from a calibration-bead sample prior to imaging. See ref.^41^ for further details.

#### 3D localization (yeast DNA loci)

We imaged fluorescently labeled DNA loci in fixed *Saccharomyces cerevisiae* yeast cells. The cells were tagged with mCherry, generating red fluorescent spots at the *POA1* DNA loci (chromosome II); each cell exhibits a single fluorescent spot^38^. Imaging used a 4f extension to the optical system (equipped with an oil-immersion objective, NA = 1.49, 100× magnification), incorporating a Tetrapod phase mask designed for an axial range of 3 µm. The PSF model was retrieved using VIPR^48^ from a calibration bead. Two datasets were acquired from different fields of view and experimental stages: one with high SNR and relatively sparse cells, and one with low SNR and higher density (including naturally denser FOVs resulting from cell aggregation, acquired later after some photobleaching). The dense, low-SNR dataset served as a challenging dataset for DS3D localization; the high-SNR dataset served as the test set for the localization-precision estimate reported in the main text.

#### 3D localization (bacteria)

This dataset, from Karempudi et al.^49^, consists of 3D imaging of chromosomal loci labeled with YFP in live *E. coli*, using an oil-immersion objective (NA = 1.45, 100× magnification) with a cylindrical lens to introduce astigmatism for depth encoding. See Supplementary Sec. A for further details.

## Code availability

The code is available online at: https://github.com/ofrigold/pilpel

## Acknowledgments

We thank Prof. Johan Elf and Axel Truedson for helpful discussions. We also thank Dr Elisa Dultz and Prof. Karsten Weis for providing us with the yeast strains used in the yeast cells experiments.

This research was supported by the ISRAEL SCIENCE FOUNDATION (grant No. 450/18). OG was supported in part by VATAT Scholarship Program for PhD Students in Data Science.

## Footnotes

* In our work, we use the improved DLPv2^36^ as our base model and refer to it simply as DLP.

## References

1. Betzig, E. et al. Imaging Intracellular Fluorescent Proteins at Nanometer Resolution. Science 313, 1642–1645 (2006).

2. Rust, M. J., Bates, M. & Zhuang, X. Sub-diffraction-limit imaging by stochastic optical reconstruction microscopy (STORM). Nature Methods 3, 793–796 (2006).

3. Sharonov, A. & Hochstrasser, R. M. Wide-field subdiffraction imaging by accumulated binding of diffusing probes. Proceedings of the National Academy of Sciences 103, 18911– 18916 (2006).

4. Sahl, S. J., Hell, S. W. & Jakobs, S. Fluorescence nanoscopy in cell biology. Nature Reviews Molecular Cell Biology 18, 685–701 (2017).

5. Shen, H. et al. Single Particle Tracking: From Theory to Biophysical Applications. Chemical Reviews 117, 7331–7376 (2017).

6. Huang, B., Wang, W., Bates, M. & Zhuang, X. Three-Dimensional Super-Resolution Imaging by Stochastic Optical Reconstruction Microscopy. Science 319, 810–813 (2008).

7. Pavani, S. R. P. et al. Three-dimensional, single-molecule fluorescence imaging beyond the diffraction limit by using a double-helix point spread function. Proceedings of the National Academy of Sciences 106, 2995–2999 (2009).

8. Shechtman, Y., Sahl, S. J., Backer, A. S. & Moerner, W. E. Optimal Point Spread Function Design for 3D Imaging. Physical Review Letters 113, 133902 (2014).

9. Shechtman, Y., Weiss, L. E., Backer, A. S., Sahl, S. J. & Moerner, W. E. Precise Three-Dimensional Scan-Free Multiple-Particle Tracking over Large Axial Ranges with Tetrapod Point Spread Functions. Nano Letters 15, 4194–4199 (2015).

10. Shechtman, Y., Weiss, L. E., Backer, A. S., Lee, M. Y. & Moerner, W. E. Multicolour localization microscopy by point-spread-function engineering. Nature Photonics 10, 590– 594 (2016).

11. Smith, C., Huisman, M., Siemons, M., Grünwald, D. & Stallinga, S. Simultaneous measurement of emission color and 3D position of single molecules. Optics Express 24, 4996 (2016).

12. Backer, A. S. & Moerner, W. E. Single-molecule orientation measurements with a quadrated pupil. Optics Letters 38, 1521–1523 (2013).

13. Hulleman, C. N. et al. Simultaneous orientation and 3D localization microscopy with a Vortex point spread function. Nature Communications 12, 5934 (2021).

14. Goldenberg, O., Ferdman, B., Nehme, E., Ezra, Y. S. & Shechtman, Y. Learning Optimal Multicolor PSF Design for 3D Pairwise Distance Estimation. Intelligent Computing 2022, 0004 (2022).

15. Lelek, M. et al. Single-molecule localization microscopy. Nature Reviews Methods Primers 1, 39 (2021).

16. Goldenberg, O., Xiao, D. & Shechtman, Y. Deep Learning for Localization Microscopy in 2D and3D. Accounts of Chemical Research 59, 2450–2461 (2026).

17. Nehme, E., Weiss, L. E., Michaeli, T. & Shechtman, Y. Deep-STORM: super-resolution single-molecule microscopy by deep learning. Optica 5, 458 (2018).

18. Nehme, E. et al. DeepSTORM3D: dense 3D localization microscopy and PSF design by deep learning. Nature Methods 17, 734–740 (2020).

19. Speiser, A. et al. Deep learning enables fast and dense single-molecule localization with high accuracy. Nature Methods 18, 1082–1090 (2021).

20. Hershko, E., Weiss, L. E., Michaeli, T. & Shechtman, Y. Multicolor localization microscopy and point-spread-function engineering by deep learning. Optics Express 27, 6158 (2019).

21. Jouchet, P., Roy, A. R. & Moerner, W. E. Combining deep learning approaches and point spread function engineering for simultaneous 3D position and 3D orientation measurements of fluorescent single molecules. Opt Commun 542, 129589 (2023).

22. Wu, T., Lu, P., Rahman, Md. A., Li, X. & Lew, M. D. Deep-SMOLM: deep learning resolves the 3D orientations and 2D positions of overlapping single molecules with optimal nanoscale resolution. Opt. Express 30, 36761–36780 (2022).

23. Zhang, P. et al. Analyzing complex single-molecule emission patterns with deep learning. Nature Methods 15, 913–916 (2018).

24. Nehme, E. et al. Learning Optimal Wavefront Shaping for Multi-Channel Imaging. IEEE Transactions on Pattern Analysis and Machine Intelligence 43, 2179–2192 (2021).

25. Sage, D. et al. Super-resolution fight club: assessment of 2D and 3D single-molecule localization microscopy software. Nature Methods 16, 387–395 (2019).

26. Archit, A. et al. Segment Anything for Microscopy. Nature Methods 22, 579–591 (2025).

27. Gupta, A. et al. SubCell: Vision foundation models for microscopy capture single-cell biology. (2024) doi:10.1101/2024.12.06.627299.

28. Ma, C., Tan, W., He, R. & Yan, B. Pretraining a foundation model for generalizable fluorescence microscopy-based image restoration. Nature Methods 21, 1558–1567 (2024).

29. Saguy, A. et al. This Microtubule Does Not Exist: Super-Resolution Microscopy Image Generation by a Diffusion Model. Small Methods 9, 2400672 (2025).

30. Baniukiewicz, P., Lutton, E. J., Collier, S. & Bretschneider, T. Generative Adversarial Networks for Augmenting Training Data of Microscopic Cell Images. Frontiers in Computer Science 1, 10 (2019).

31. Goodfellow, I. et al. Generative Adversarial Nets. in Advances in Neural Information Processing Systems vol. 27 (Curran Associates, Inc., 2014).

32. Isola, P., Zhu, J.-Y., Zhou, T. & Efros, A. A. Image-To-Image Translation With Conditional Adversarial Networks. in Proceedings of the IEEE Conference on Computer Vision and Pattern Recognition 1125–1134 (2017).

33. Zhu, J.-Y., Park, T., Isola, P. & Efros, A. A. Unpaired Image-to-Image Translation using Cycle-Consistent Adversarial Networks. in Proceedings of the IEEE international conference on computer vision 2223–2232 (2017).

34. Truedson, A., Gras, K. & Elf, J. The 3D point spread function can be autoencoded in latent space. bioRxiv https://doi.org/10.1101/2024.10.14.618231 (2024) doi:10.1101/2024.10.14.618231.

35. Daniel, T. & Tamar, A. Unsupervised Image Representation Learning with Deep Latent Particles. in International Conference on Machine Learning 4644–4665 (PMLR, 2022).

36. Daniel, T. & Tamar, A. DDLP: Unsupervised Object-centric Video Prediction with Deep Dynamic Latent Particles. Transactions on Machine Learning Research (2024).

37. Opatovski, N. et al. Multiplexed PSF engineering for three-dimensional multicolor particle tracking. Nano Letters 21, 5888–5895 (2021).

38. Shalev Ezra, Y., et al. High-throughput DNA repair monitoring in Saccharomyces cerevisiae suggests SSB-and DSB-induced chromatin reconfiguration. Scientific Reports 15, 32302 (2025).

39. Saguy, A. et al. DBlink: dynamic localization microscopy in super spatiotemporal resolution via deep learning. Nat Methods 20, 1939–1948 (2023).

40. Ovesný, M., Křížek, P., Borkovec, J., Švindrych, Z. & Hagen, G. M. ThunderSTORM: a comprehensive ImageJ plug-in for PALM and STORM data analysis and super-resolution imaging. Bioinformatics 30, 2389–2390 (2014).

41. Orange kedem, R. et al. Near index matching enables solid diffractive optical element fabrication via additive manufacturing. Light: Science & Applications 12, 222 (2023).

42. Saguy, A. et al. One-click image reconstruction in single-molecule localization microscopy via deep learning. bioRxiv https://doi.org/10.1101/2025.04.13.648574 (2025) doi:10.1101/2025.04.13.648574.

43. Backlund, M. P., Joyner, R., Weis, K. & Moerner, W. E. Correlations of three-dimensional motion of chromosomal loci in yeast revealed by the double-helix point spread function microscope. Molecular Biology of the Cell 25, 3619–3629 (2014).

44. Liu, S. et al. Universal inverse modeling of point spread functions for SMLM localization and microscope characterization. Nature Methods 21, 1082–1093 (2024).

45. Kingma, D. P. & Welling, M. Auto-Encoding Variational Bayes. (2022) doi:10.48550/arXiv.1312.6114.

46. Haramati, D., Daniel, T. & Tamar, A. Entity-Centric Reinforcement Learning for Object Manipulation from Pixels. in The Twelfth International Conference on Learning Representations (2024).

47. Qi, C., Haramati, D., Daniel, T., Tamar, A. & Zhang, A. EC-Diffuser: Multi-Object Manipulation via Entity-Centric Behavior Generation. arXiv preprint arXiv:2412.18907 (2024).

48. Ferdman, B. et al. VIPR: vectorial implementation of phase retrieval for fast and accurate microscopic pixel-wise pupil estimation. Optics Express 28, 10179 (2020).

49. Karempudi, P. et al. Three-dimensional localization and tracking of chromosomal loci throughout the Escherichia coli cell cycle. Communications Biology 7, 1443 (2024).

