## Supplementary Material for "Self-supervised generation of realistic training data enables nanoscale localization in challenging conditions"

### A. 3D localization in bacteria with astigmatism PSF

To further validate the versatility of our method across different biological systems and PSF-engineering approaches, we applied PILPEL to 3D localization data from bacteria using astigmatism-based depth encoding. This dataset, published by Karempudi et al.[1], involves imaging of fluorescently labeled chromosomal loci in live *Escherichia coli* cells. In this experiment, three chromosomal loci were labeled with yellow fluorescent protein (YFP) for imaging. The cells were imaged using a 100x/1.45NA oil immersion objective in a widefield microscopy setup equipped with a cylindrical lens to introduce astigmatism for 3D localization (see Ref[1] for details). Similar to the Tetrapod PSF used in previous experiments, astigmatism also encodes axial position, but through the ellipticity and orientation of the PSF, which depend on the distance from the focal plane. The astigmatic PSF was experimentally calibrated using fluorescent beads and phase retrieval via VIPR[2]. Cells were grown and imaged in a microfluidic device, with each field of view containing 20 growth channels.

Supplementary Figure 1a illustrates PILPEL's learned decomposition of the bacterial imaging data. The first row shows five representative measured images, each containing multiple bacterial cells with visible fluorescent loci. The second row displays PILPEL's full reconstructions. The remaining rows show the PSFs and the non-PSF components. Notably, the background components reveal a distinct, structured signal corresponding to the autofluorescence background of the cells in the microfluidic channels, learned automatically by the model.

Using the PILPEL-generated labeled dataset, we trained DS3D[3] for 3D localization on the astigmatic data. Supplementary Figure 1b shows an example of the 3D localization rendering, displayed as green-on-white overlap. To assess localization accuracy in the absence of absolute ground truth, we compared the detected positions to the 3D localizations produced by the FD-DeepLoc pipeline reported in Ref[1]; observing the same hallmark spatial patterns: our reconstructions reproduced the ring-shaped yz-plane distribution of the *oriC* locus and the expected long and short-axis localizations (Supplementary Figure 1c). Quantitatively, the mean absolute error between our localizations and those from the original paper for the *oriC* data is ~ 17 nm in X and Y axes and ~ 58 nm in Z axis, over 128K localizations.

This bacterial localization experiment is significant for several reasons. First, it demonstrates again that our approach readily adapts to different PSF engineering strategies - simply by replacing the physical PSF model in the decoder, without any other architectural changes. Second, this biological system exhibits different background patterns and autofluorescence characteristics compared to the yeast cells. Notably, in order to train a localization network on their data, the original work[1] required a substantial simulation pipeline, involving cell segmentation masks of phase-contrast images, extraction of intensity profiles, and careful modeling of cell-specific background fluorescence patterns to create training data resembling the real measurements. In contrast, our PILPEL model learns these complex background characteristics and noise patterns directly from the experimental data without any manual intervention or explicit modeling of the bacterial cell morphology.

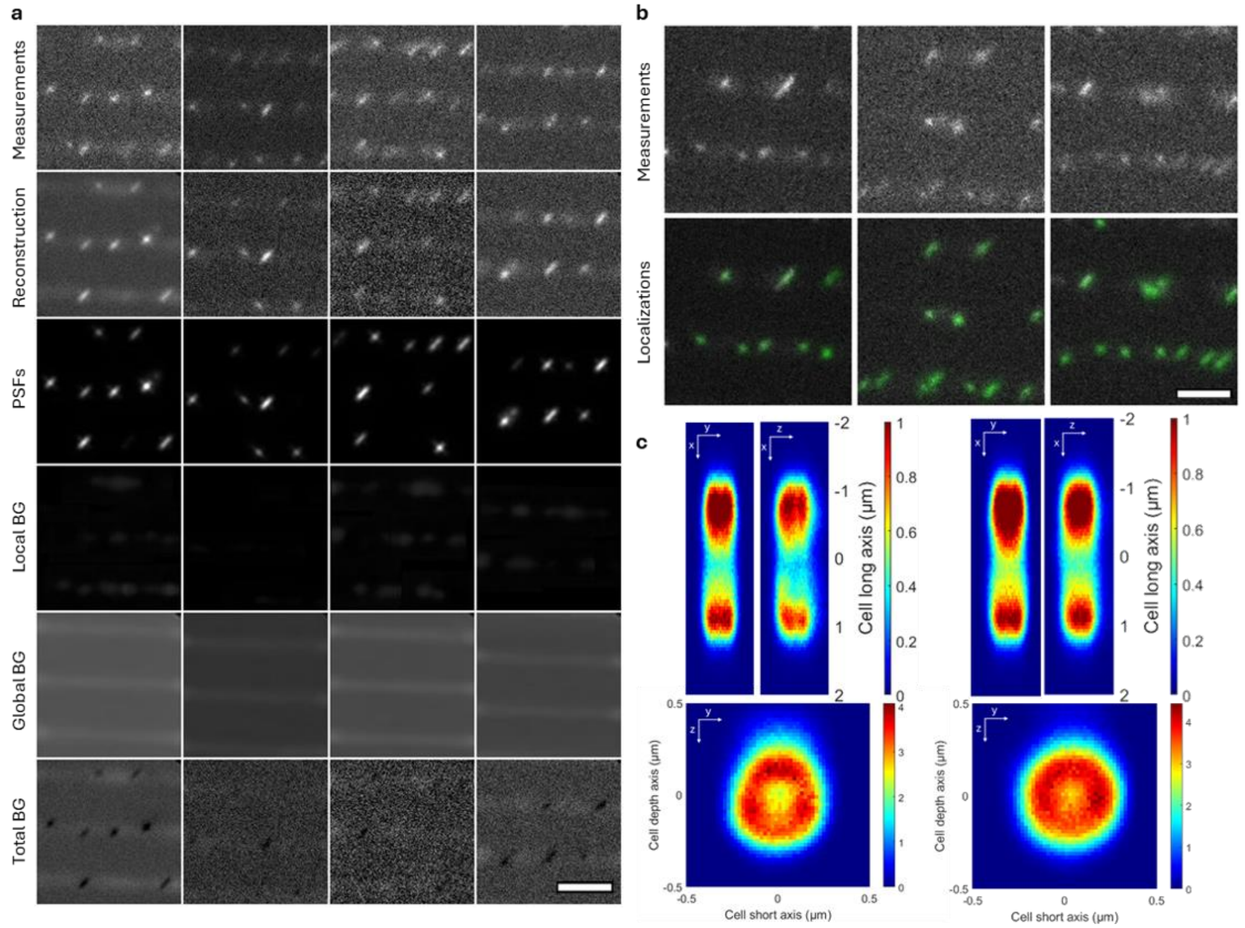


*Supplementary Figure 1: 3D localization of loci in E. coli. (a) PILPEL output components. The first row shows five measured images; the second row shows PILPEL's reconstruction; the remaining rows show the reconstruction components. The Local BG and Global BG rows display the distinct background corresponding to the cells in the microfluidic channels, learned by the model. (b) 3D localization rendering by DS3D (DeepSTORM3D) trained on PILPEL-generated data. (c) 3D spatial distribution of the oriC locus in cell-internal coordinates. The data are two-dimensional histograms of localized emitter positions, pooled from time-lapse imaging across all detected cell areas. Results on the left are from the original paper, and results on the right are ours, demonstrating a highly similar localization pattern. The top panel shows the distribution along the cell's long (x) and short (y) axes, and the bottom panel shows the radial distribution in the short (y) and depth (z) axes (yz-plane). Scale bar, 2 µm.*

### B. Controlled simulation for performance assessment

To generate diverse and challenging noise patterns for the `PerlinData` test set, our implementation uses Perlin noise scale parameters ranging from 0.5 to 1.7 and intensity values from 5 to 14, which control the spatial frequency and amplitude of the noise, respectively. These parameters were chosen to produce diverse patterns—noisy enough to challenge localization accuracy, but not so severe as to obscure the PSF signal. The results presented here provide a detailed analysis of the performance gains referenced in the main text.

The histogram in Supplementary Figure 2a compares the 3D localization error distributions for DS3D trained on the "General bg" versus the "PILPEL" dataset. Visually, the PILPEL-trained model's distribution (green) is clearly shifted toward lower errors, with a higher peak and a smaller high-error tail than the original model (blue). This improvement is quantified by the mean absolute error, which dropped from 0.071 μm for the original model to 0.053 μm for the PILPEL-trained model.

The table in Supplementary Figure 2b further details these robustness gains, breaking down the performance ($\Delta$RMSE and $\Delta$Jacc) when comparing the PILPEL-trained model against two different baselines: the standard "General bg" simulator and the "Yeast bg" simulator (which included manual prior knowledge). When compared to the "General bg" baseline, the PILPEL model improved $\Delta$RMSE by $-19$ nm ($-28\%$) and $\Delta$Jacc by $+0.47$ on the `Perlin` test set. On the more difficult `Perlin Low SNR` set, the error reduction was $-21$ nm ($-26\%$) with a $\Delta$Jacc gain of $+0.21$. Critically, PILPEL also outperformed the model trained with manually added prior knowledge ("Yeast bg"). Against this stronger baseline, PILPEL was still better by $-12$ nm ($-16\%$) $\Delta$RMSE and $+0.22$ $\Delta$Jacc on the challenging `Perlin Low SNR` set. Lateral and axial RMSE results are presented in Supplementary Figure 3. These results confirm that PILPEL's learned generative approach is more effective than training on manually-designed synthetic datasets, with the large Jaccard index gains demonstrating a higher rate of correct emitter detection.

a

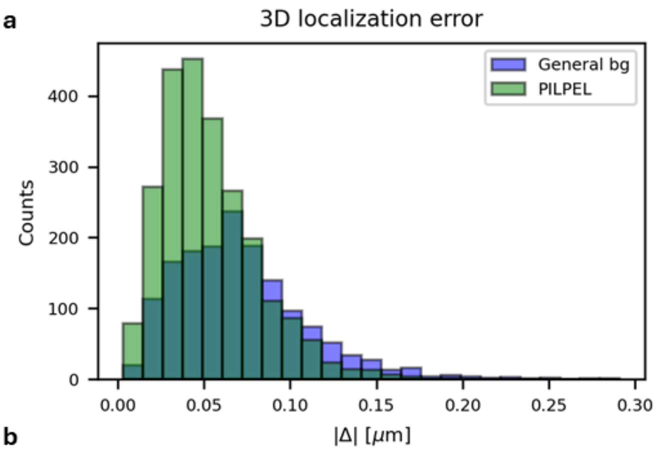


b

| | PILPEL vs. General bg | | | PILPEL vs. Yeast bg | | |
|---|---|---|---|---|---|---|
| Test set | ΔRMSE | | Δ Jacc | ΔRMSE | | Δ Jacc |
| Perlin | -19 [nm] | **-28 [%]** | **+0.47** | -5 [nm] | -9 [%] | +0.40 |
| Perlin Low SNR | -21 [nm] | **-26 [%]** | **+0.21** | -12 [nm] | -16 [%] | +0.22 |

*Supplementary Figure 2: Robustness experiment on simulations– DS3D localization performance. (a) 3D localization error comparison: DS3D trained on the generated dataset vs. DS3D trained on the standard simulated dataset. Mean absolute error for the generated model is 0.053 μm, compared to 0.071 μm for the original. (b) Improvement of training on generated data. Localization RMSE and Jaccard index for DS3D trained on the generated dataset vs. training on yeast-BG and original-DS3D datasets.*

a

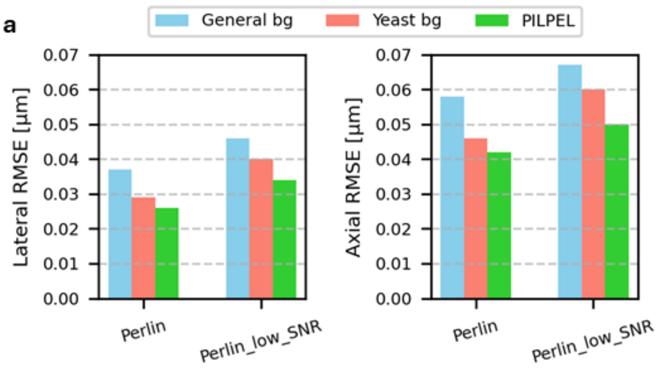


b

| | PILPEL vs. General bg | | | | PILPEL vs. Yeast bg | | | |
|---|---|---|---|---|---|---|---|---|
| Test set | ΔLateralRMSE | | ΔAxialRMSE | | ΔLateralRMSE | | ΔAxialRMSE | |
| Perlin | -11 [nm] | -29 [%] | -16 [nm] | -27 [%] | -3 [nm] | -10 [%] | -4 [nm] | -8 [%] |
| Perlin Low SNR | -12 [nm] | -26 [%] | -17 [nm] | -25 [%] | -6 [nm] | -15 [%] | -10 [nm] | -17 [%] |

*Supplementary Figure 3: Lateral and axial localization accuracy comparison. Quantitative evaluation of the same experiment shown in Supplementary Figure 2, with the RMSE separated into lateral ($x, y$) and axial ($z$) components. Results are reported for all evaluated methods and datasets, demonstrating consistent improvement of our approach in both dimensions.*

### C. Experimental evaluation of super-resolution reconstructions

To evaluate PILPEL's improvement on the experimental data lacking an absolute ground truth, we analyzed the localization results and super-resolution reconstructions of the 2D and 3D STORM experiments, applying complementary analyses to the two resulting datasets: standard DS/DS3D localizations and PILPEL-trained localizations. PILPEL detects more localizations than the original network on the same frames, and qualitative comparisons demonstrate that PILPEL produces a denser, fuller, and more continuous reconstruction; the analyses below verify that this increase is mostly enhancing true biological structures.

First, to validate that the increase in localizations yields a more complete structural representation without introducing artifacts, we performed a line-profile analysis on the 2D STORM data of microtubules, showing that the signal increase produced by PILPEL is spatially concentrated at the microtubule filaments. Across three microtubule ROIs, we compared width-averaged intensity profiles on the filament ("center") and in two adjacent structure-free regions ("left"/"right") for DS and PILPEL (Supplementary Figure 4a). On the microtubule, PILPEL raised the mean profile intensity by 0.039, 0.087, and 0.117 a.u. over DS across the three ROIs, whereas in the empty regions the increase stayed at the background floor (≤0.006 a.u.). Thus, while false positives can occur, the extra localizations PILPEL recovers are predominantly associated with genuine microtubule signal.

Similarly on the 3D STORM of mitochondria, we performed density analysis on 20 regions of interest (ROIs) containing distinct mitochondrial structures and 20 control ROIs situated in empty background regions (Supplementary Figure 5a). We defined 20 ROIs on mitochondrial tubules, each a rectangular box in *x–y* restricted to the axial slab containing that tubule. Each ROI was translated laterally onto an adjacent structure-free region, preserving box dimensions and *z*-slab. This yields 20 matched pairs differing only in the presence of a mitochondrion—identical volumes, axial position, and confidence threshold (40) for both methods (Supplementary Figure 5b). Densities are per unit volume (loc/µm³) with total volume of 11.18 µm³ per group. The added density is 635.5 loc/µm³ on structure and 4.11 loc/µm³ off structure (see Table 1), and the median nearest-neighbor distance on the structure-areas was finer in all 20 ROIs (Supplementary Figure 5e).

*Table 1: Localization counts and densities in structure and background ROIs for DS3D and PILPEL-trained networks. Pooled over all ROIs, n = 20 matched ROI pairs, total volume 11.18 µm³ per group.*

| | **DS3D** | **PILPEL** |
|---|---|---|
| On-structure localizations | 3,329 (297.9 loc/µm³) | 10,432 (933.4 loc/µm³) |
| Off-structure localizations | 5 (0.45 loc/µm³) | 51 (4.56 loc/µm³) |

In a second analysis for biological structure validation, we measured the apparent filament width from cross-sectional profiles of isolated microtubules (Supplementary Figure 4b). Gaussian fits yielded a mean FWHM of 66.4 ± 5.9 nm for DS and 68.1 ± 4.6 nm for PILPEL (consistent with the apparent width expected for antibody-labeled microtubules[4]), with no significant difference between the methods. Critically, PILPEL's higher detection rate adds signal along the microtubules without artificially broadening them. In the 3D data, an axial cross-section through a mitochondrial tubule resolves the hollow cross-section, matching the expected mitochondrial geometry (Supplementary Figure 5c and d).

Third, we evaluated the reconstructions empirically using Fourier Ring Correlation (FRC)[5] on independent even/odd frame splits of the microtubule dataset (Supplementary Figure 4c). Stochastic hallucinated detections would not correlate between the two halves and would lower the measured resolution. At the full field-of-view (all counts), the methods yielded comparable global resolutions (DS 63 nm, PILPEL 60 nm), with PILPEL carrying approximately twice the localizations. In addition to the full field, we repeated the FRC analysis in fixed 5 µm tiles across the field of view (n = 32; Supplementary Figure 4d). Consistent with the global result, PILPEL was comparable to DS per tile, with a modest median improvement of ~4 nm.

Together, these results indicate that PILPEL's additional localizations are consistent with genuine biological structure and mostly confined to regions occupied by structure.

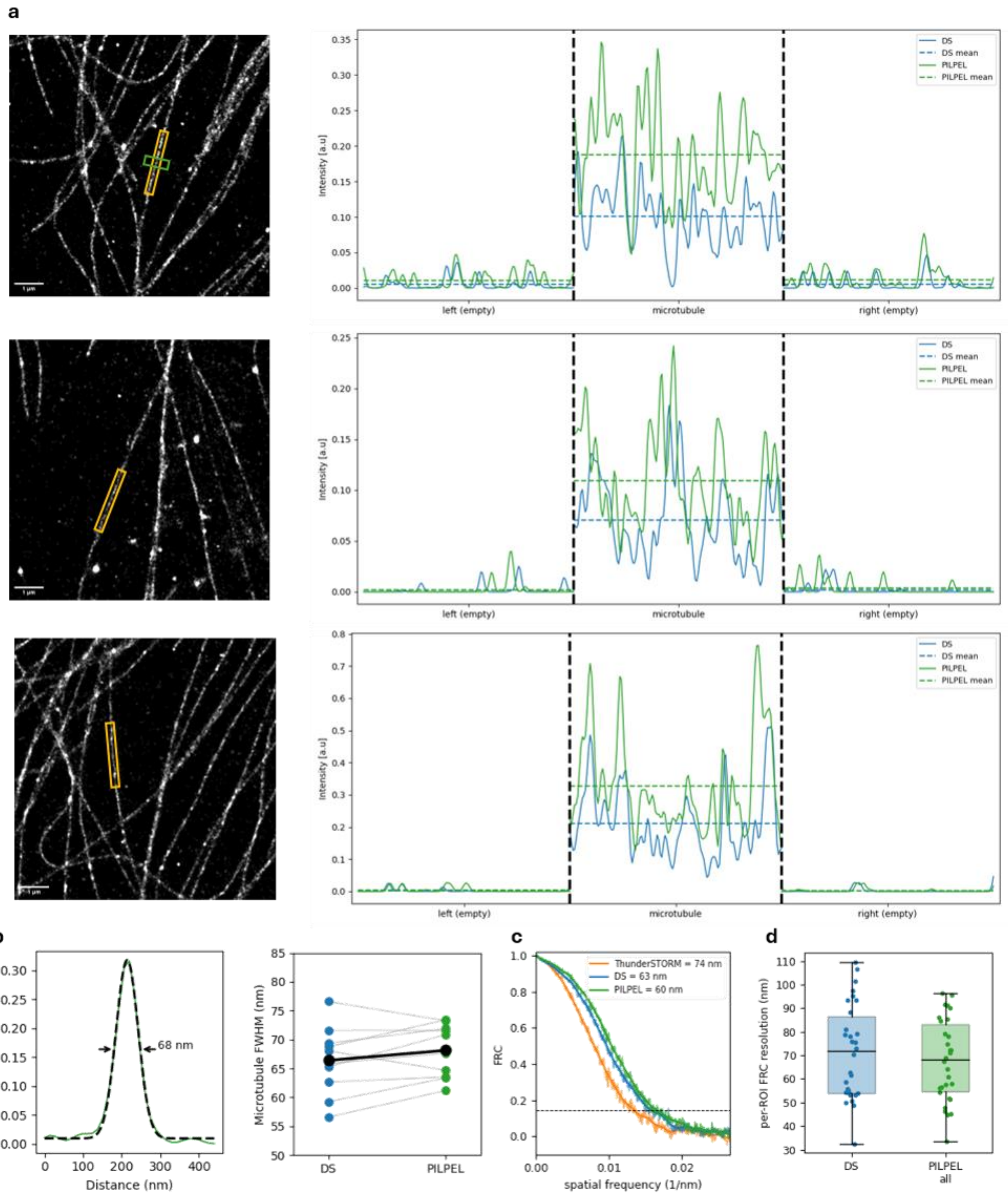


*Supplementary Figure 4: Validation of 2D STORM reconstructions. (a) Super-resolution reconstructions of microtubule filaments and corresponding intensity line profiles across selected regions of interest (yellow ROIs). The profiles compare the original DeepSTORM (DS) localizations with PILPEL-trained localizations, showing that the signal increase is concentrated at the genuine microtubule structure (center zone) and remains at the near-zero background floor in adjacent structure-free regions (left/right zones).(b) Validation of apparent filament width. Left: A representative Gaussian cross-section fit of an isolated microtubule*

*(green box, top row of (a)). Right: Paired comparison of FWHM measurements across n=10 ROIs. PILPEL's higher detection rate preserves the apparent filament width without artificial broadening (mean FWHM: DS 66.4 ± 5.9 nm, PILPEL 68.1 ± 4.6 nm). (c) Global Fourier Ring Correlation (FRC) evaluating resolution on independent even/odd frame splits, where resolution is the inverse of the spatial frequency at which the FRC curve crosses the 1/7 threshold (dashed line). ThunderSTORM: 74 nm, DS: 63 nm, PILPEL: 60 nm. (d) Per-ROI FRC resolution across fixed 5 µm tiled regions (n = 32). PILPEL is comparable to DS with slight improvement while detecting 1.96× more localizations.*

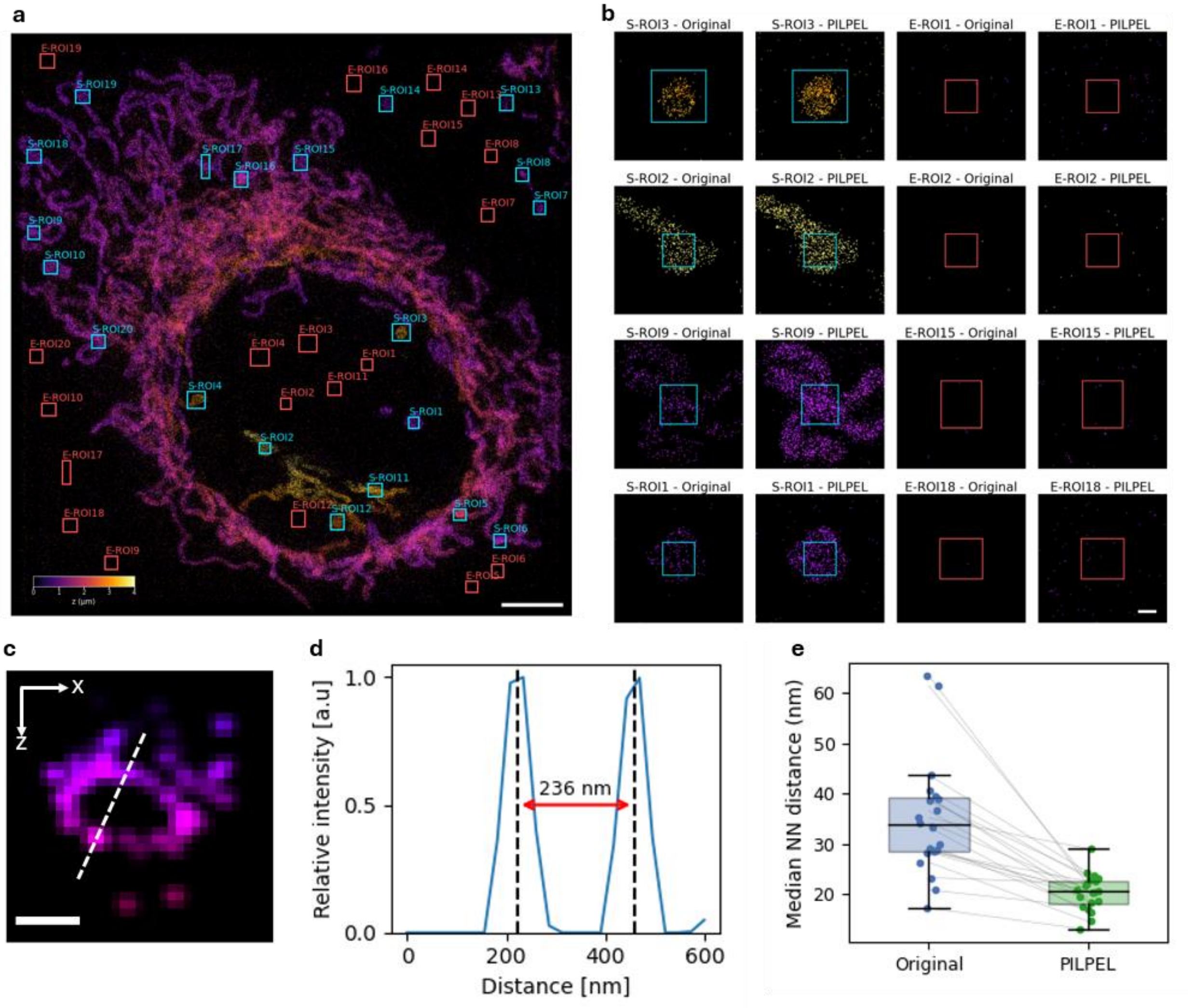


*Supplementary Figure 5: Volumetric density and structural validation of 3D STORM mitochondria reconstructions. (a) Full field-of-view 3D STORM reconstruction color-coded by axial depth (z). For density evaluation, 20 on-structure regions of interest (S-ROIs, cyan boxes) were defined on mitochondrial tubules, and 20 volume-matched off-structure background regions (E-ROIs, red boxes) were selected in close empty areas. Scale bar: 5 µm. (b) Representative x-y projections of on-structure and off-structure ROIs from (a), comparing the Original (DS3D) and PILPEL-trained reconstructions. Scale bar: 0.5 µm (c) Axial cross-section (x-z plane) of a PILPEL-reconstructed mitochondrial tubule, successfully resolving the expected hollow biological structure. Scale bar: 0.2 µm. (d) Intensity profile measured along the dashed*

*line in (c). The distance between the two peaks yields an apparent tubule diameter of 236 nm. (e) Paired comparison of the median nearest-neighbor (NN) distance across the 20 on-structure ROIs. PILPEL consistently yields a finer median NN distance than the baseline across all measured regions.*

### D. Robustness to PSF aberrations

We tested robustness to PSF mismatch by perturbing the VIPR phase mask with a fixed random Zernike combination (astigmatism, coma, spherical) at RMS wavefront error amplitudes ranging from a perfect match (0 mλ) to severe mismatch (150 mλ). PILPEL was retrained from scratch on PerlinData with each miscalibrated prior (used in both the initial proposal module and the physical decoder, as in the usual pipeline), then a fresh DS3D was trained on its generated images. On the test set Jaccard remains high within the diffraction-limited regime (J = 0.9, RMSE = 41 nm at 30 mλ; J = 0.85, RMSE = 60 nm at 60 mλ), degrades sharply once mismatch exceeds the Maréchal threshold (≈71 mλ), reaching J = 0.41, RMSE = 80 nm at 100 mλ, and collapses at 150 mλ (J = 0.08) under extreme miscalibration (see Supplementary Figure 6). These results quantify the prior-PSF dependence noted in the limitations section: PILPEL tolerates miscalibration within the diffraction-limited regime, while larger mismatch beyond the Maréchal threshold degrades performance. Future work could remedy this limitation by introducing an additional latent variable to learn and model pupil function aberrations.

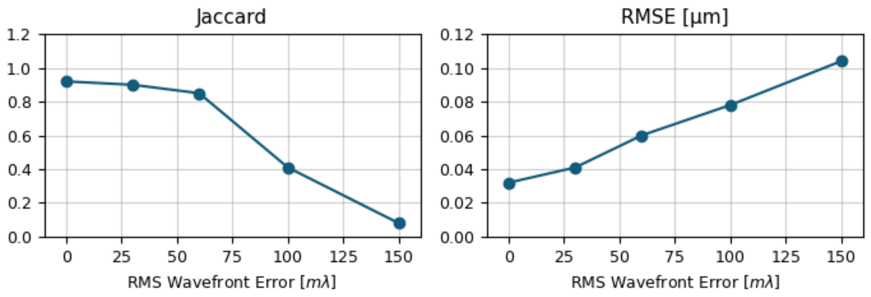


*Supplementary Figure 6: Aberration robustness. Downstream localization performance (Jaccard and RMSE) of DS3D trained on PILPEL-generated datasets, as the PSF model is increasingly miscalibrated.*

### E. Ablation Study

To quantify the contribution of each component in the PILPEL framework, we evaluated the localization performance on the `PerlinData` test set under different configurations. Supplementary Figure 7 presents the Precision-Recall curves for all variants, and Table 2 summarizes the key metrics, calculated at a working confidence threshold of 100. The results demonstrate that explicitly modeling the background is critical. Removing the **Global Background** module leads to a collapse in performance (recall < 1%). This occurs primarily because artifacts in the generated images cause a large deviation from the test set; consequently, most PSFs are drowned to zero as the local object components attempt to compensate and learn the general image structure. The **Local Background** is also important for reconstruction, and its removal degrades performance, though the degradation is less severe, likely due to the presence of the global background. Removing the **intensity-pruning** mechanism prevents the model from zeroing out spurious proposals. Since all initialized particles remain active at fixed high intensity, the generated training images are flooded with emitters at random locations, producing unrealistically dense and cluttered training data, causing recall to be capped at ~0.3 regardless of detection threshold, as seen in Supplementary Figure 7. While this configuration yields slightly higher precision (0.97) among all variants, it comes with a drastic drop in sensitivity, and the Jaccard index (0.21) correctly captures the severity of this failure. Finally, omitting the **template-matching** initialization results in unreliable particle location proposals, leading to an excess of false detections and degraded precision throughout, which reduces the Jaccard index to 0.75. The full PILPEL configuration yields the highest overall performance as measured by the Jaccard index (0.92) and the lowest localization error (RMSE 0.032 μm), confirming the necessity of all proposed modules.

*Table 2: Ablation Study Results*

| **Configuration** | **Precision** (↑) | **Recall** (↑) | **Jaccard** (↑) | **RMSE (μm)** (↓) |
|---|---|---|---|---|
| **Full PILPEL** | 0.93 | **0.99** | **0.92** | **0.032** |
| w/o Local Background | 0.81 | 0.98 | 0.80 | 0.035 |
| w/o Global Background | 0.38 | 0.005 | 0.005 | 0.054 |
| w/o intensity-pruning | **0.97** | 0.21 | 0.21 | 0.051 |
| w/o template-matching | 0.8 | 0.91 | 0.75 | 0.034 |

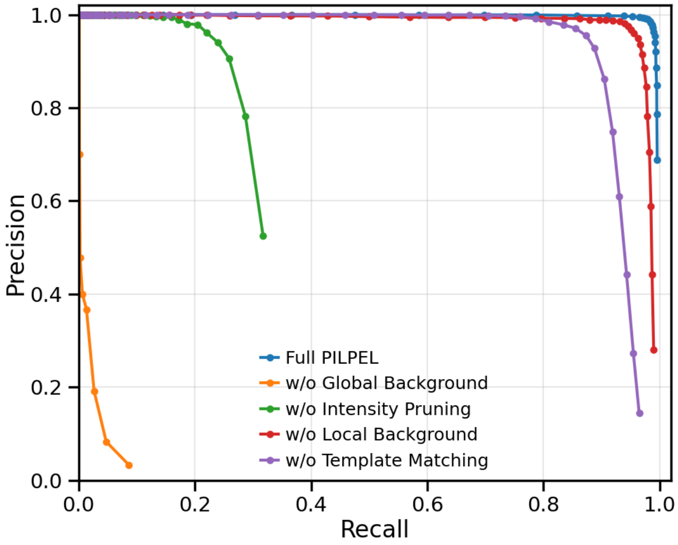


*Supplementary Figure 7: Ablation Study: Precision-Recall Curves. Performance comparison of the full PILPEL model (blue) against configurations with individual modules removed. The full model yields the most robust detection envelope.*

### F. Computational details

All PILPEL training runs were performed on a single NVIDIA Titan RTX GPU (32 GB) using PyTorch. We trained the model for 50 epochs with a batch size of 4 using Adam optimizer, and a learning rate of 1e-5. We held out 10% of each dataset for validation. The reconstruction loss was either pixel-wise MSE or a VGG-based perceptual loss[6]. Based on our empirical observations across datasets, MSE appears preferable for noisy backgrounds with minimal structure (blinking emitters), while perceptual loss better suits textured backgrounds with defined patterns (yeast, Perlin noise); however, a systematic evaluation across more datasets is required to draw definitive conclusions.

For each dataset, we performed VIPR[2] phase retrieval on calibration-bead measurements to estimate a phase mask with experiment-specific optical parameters and the expected axial range (see Supplementary Figure 10). The resulting PSF model served both as the differentiable PSF generator in the decoder and as the basis for a PSF dictionary used by the correlation-based prior for initial coordinate proposals. We discretized the PSF into 21 $z$-planes across the modeled axial range. To generate the proposals, we partitioned the image into a uniform grid of patches, computed the 3D cross-correlation of each patch with the dictionary, and re-centered the patches around the coordinates of maximum correlation. The patch size is determined by the PSF model dimensions.

Number of particles to encode was chosen per dataset (detailed in Table 3), while high initialization used for typically dense emitters, and reduced if the data was substantially sparse, trading efficiency against potential missed localizations. The latent dimension was 10 for the local-background features and 20 for the global-background encoding. Architecture, latent dimensions and all components not modified in our method followed DLP[7], using the default recommended hyper-parameters from the open-source code. Training typically converged within ~8 hours on average, depending on the dataset and image size. After training, the decoder served as a generator to synthesize 10,000 labeled images for DS3D training. PILPEL inference (i.e. data generation) was performed on the same hardware at approximately 5 frames/s, depending on image size.

Dataset-specific PILPEL configurations and training details appear in Table 3.

*Table 3: PILPEL hyperparameters per dataset*

| Parameter | Perlin | 3D STORM | Yeast | 2D STORM | Bacteria |
|---|---|---|---|---|---|
| Learning rate | 1e-4 | 1e-5 | 1e-4 | 1e-5 | 1e-5 |
| Reconstruction loss type | VGG | MSE | VGG | MSE | MSE |
| Image size (px) | 193 | 129 | 193 | 65 | 129 |
| Patch size (px) | 32 | 32 | 32 | 16 | 32 |
| Initial latent particles # | 60 | 16 | 60 | 36 | 16 |
| Training set size (images) | 1000 | 3000 | 1200 | 4200 | 4000 |
| Training time (min/epoch) | 7.70 | 6.57 | 9.72 | 11.76 | 14.46 |
| Total runtime (h) | 6.4 | 5.5 | 8.1 | 9.8 | 12 |
| Inference time (images/min) | 155 | 571 | 180 | 409 | 380 |

### G. Supplementary Figures

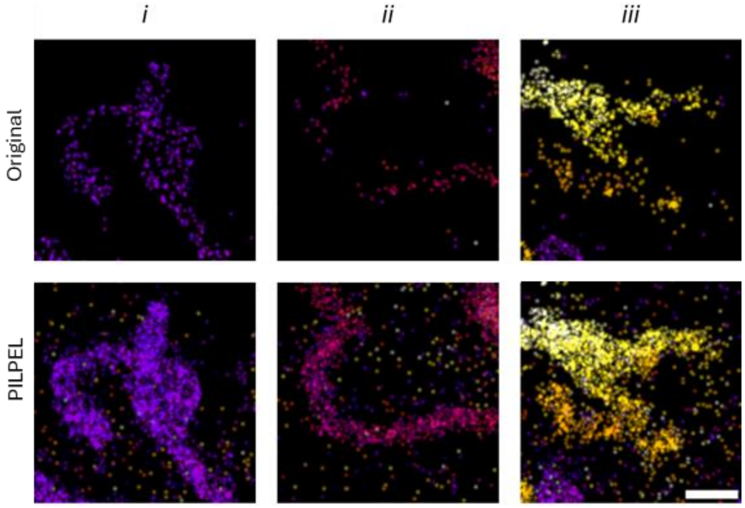


*Supplementary Figure 8: Unfiltered 3D super-resolution reconstruction of mitochondria. Magnified views of regions i, ii, and iii from Figure 5 in the main text. Top row: reconstruction using DS3D trained on standard simulated data. Bottom row: reconstruction using DS3D trained on PILPEL-generated data. As discussed in the main text, the PILPEL-trained reconstruction shown in Figure 5 was post-processed with a density filter to facilitate clearer visual comparison. Here, we show the raw, unfiltered PILPEL-trained reconstruction. As can be seen, the higher number of detections enables more complete structural recovery, which in turn allows such filtering to enhance visualization. In contrast, the standard-trained reconstruction is more sparse and applying the same filter would erase the remaining structural information. Scale bar, 1 µm.*

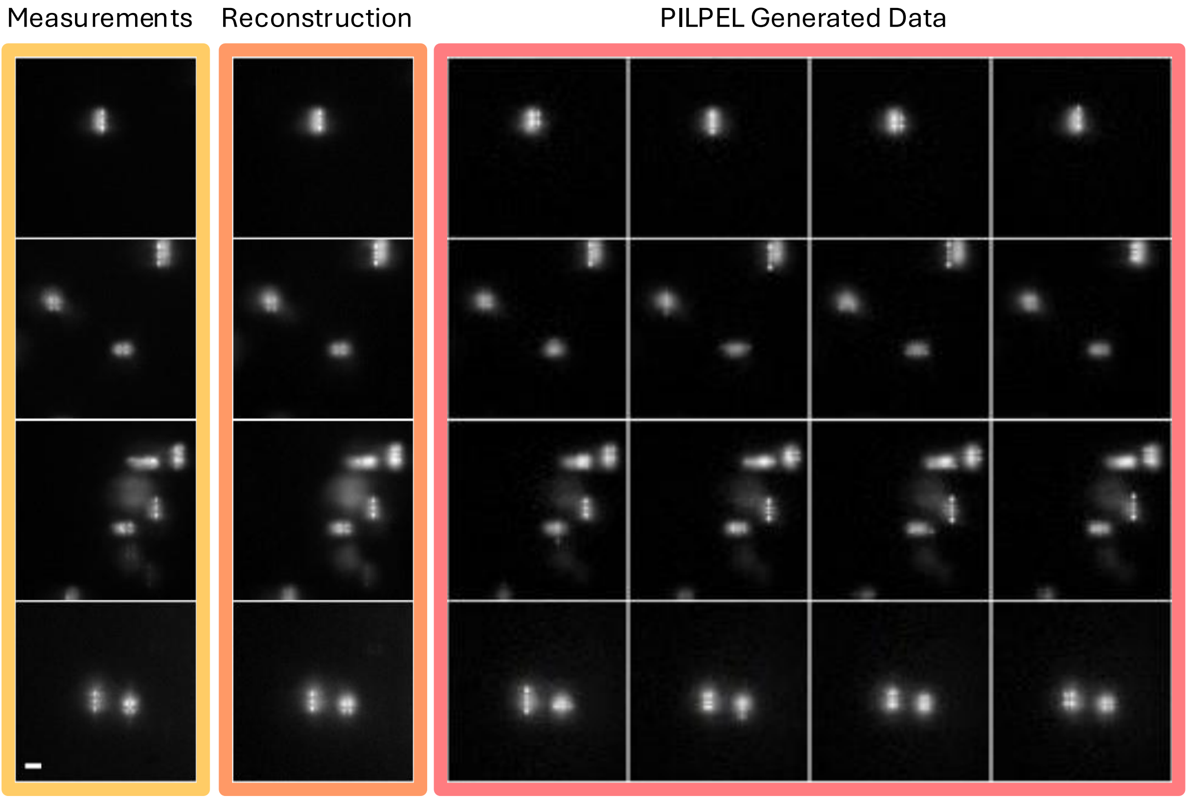


*Supplementary Figure 9: Realistic labeled training data generation. Measured images are passed through the encoder-decoder model to generate the reconstructions. To create a labeled dataset, the encoded PSF attributes are randomly perturbed, and the stochastic latent variables are resampled. This process produces realistic image variants that resemble the original measurements. Since the PSF coordinates fed into the generator are known, the resulting dataset is fully labeled and can be used to train a 3D localization network. Scale bar, 2 µm*

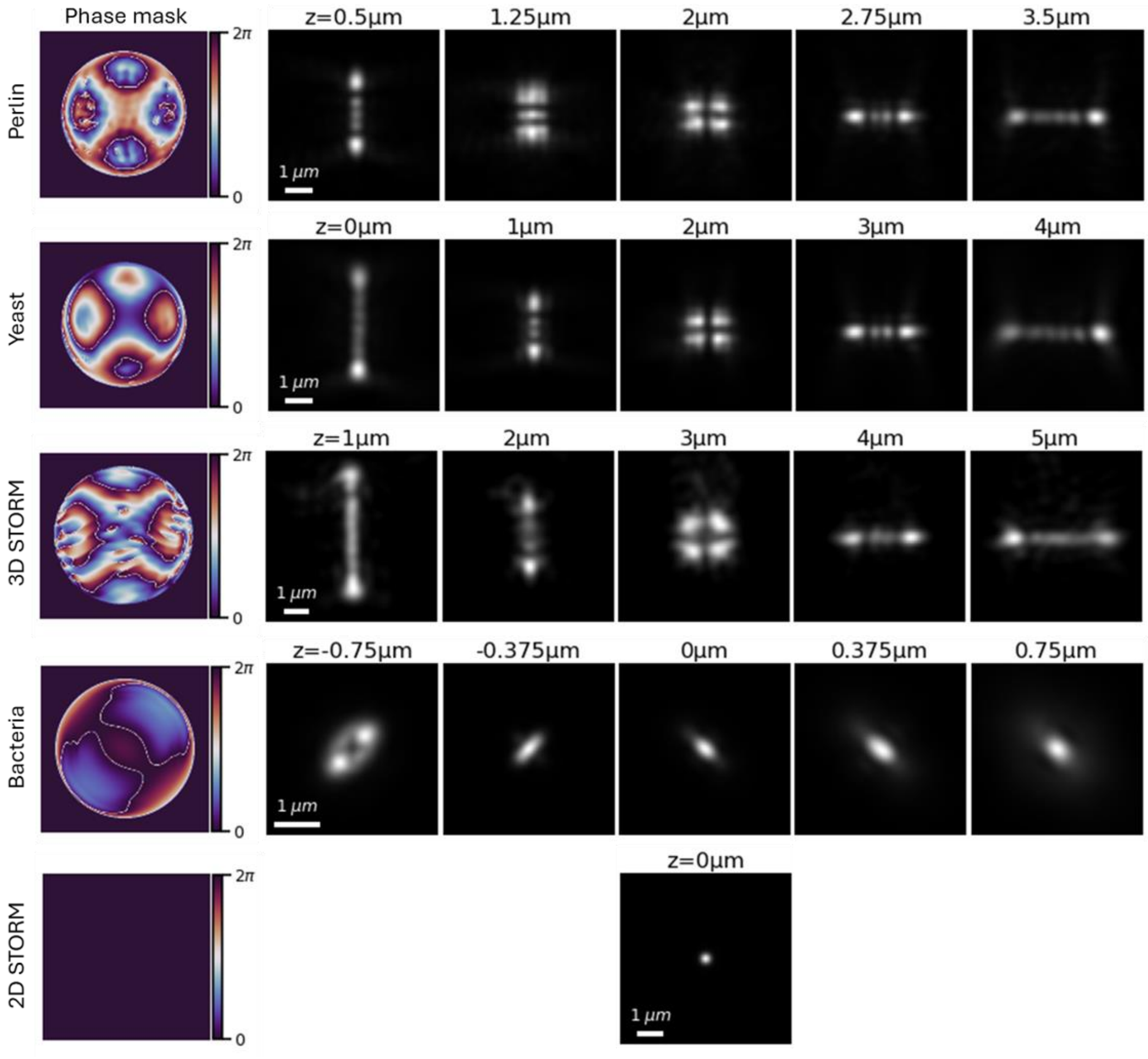


*Supplementary Figure 10: PSF models used across experiments. Phase masks retrieved with VIPR from calibration-bead measurements and their corresponding PSF models, illustrating the depth-dependent PSF shapes used for reconstruction and data generation. Shown are Tetrapod PSFs (used in the synthetic Perlin dataset, 3D STORM of mitochondria and yeast DNA loci), an astigmatic PSF for E. coli experiment, and the standard diffraction-limited PSF used in the 2D STORM experiment*

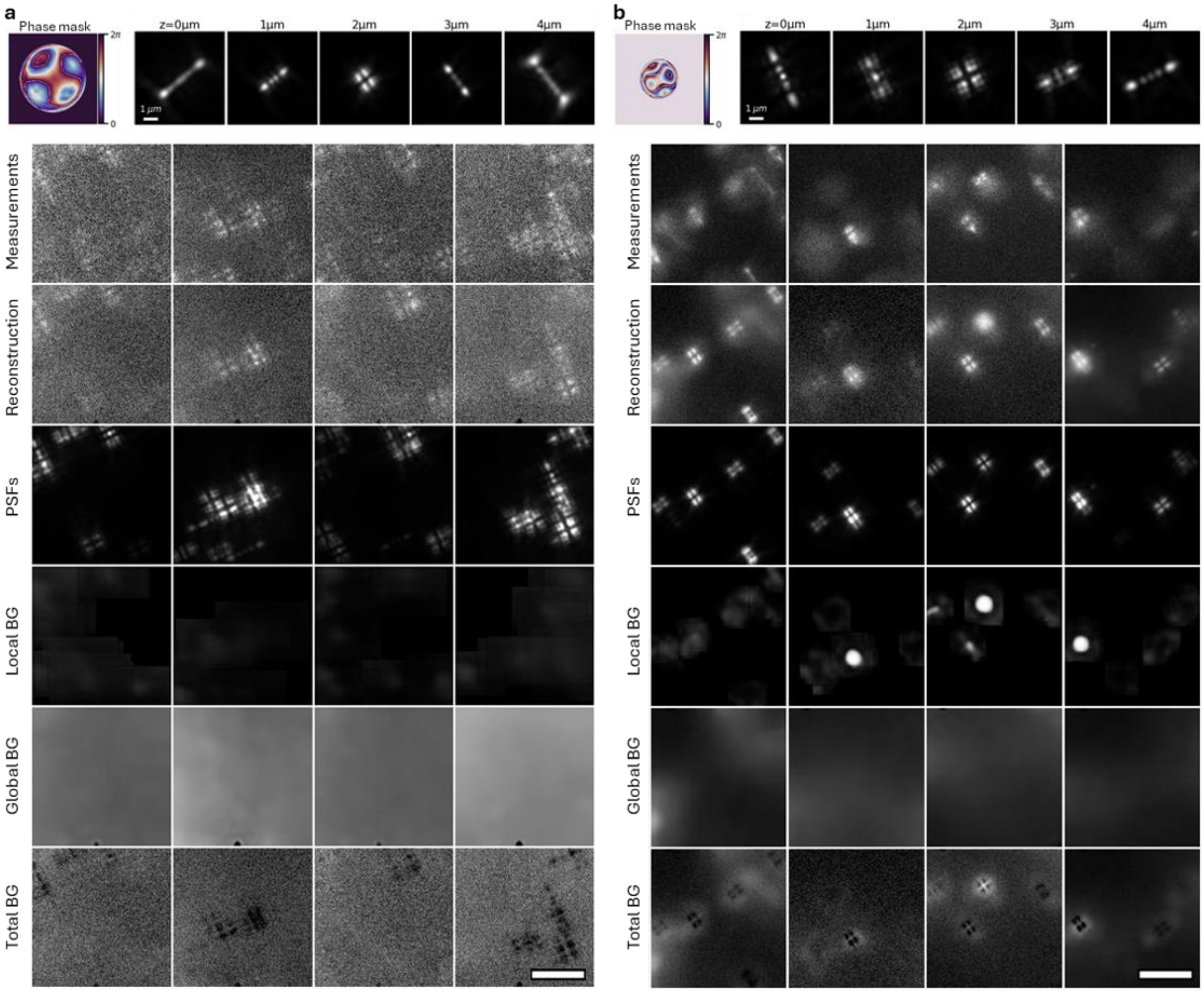


*Supplementary Figure 11: Additional PILPEL reconstructions and component decomposition of experimental data. Left: 3D STORM of mitochondria labeled with Alexa Fluor 647, and imaged with a rotated Tetrapod designed for $\lambda = 670\,nm$ and an axial range of $4\,\mu m$. Right: fluorescent DNA loci in yeast cells, labeled with eGFP and imaged with a rotated Tetrapod designed for $\lambda = 515\,nm$ and an axial range of $4\,\mu m$. Top insets show the phase masks and the calibrated PSF model retrieved with VIPR. Scale bars, 1 μm.*

### References


1. Karempudi, P. *et al.* Three-dimensional localization and tracking of chromosomal loci throughout the Escherichia coli cell cycle. *Communications Biology* **7**, 1443 (2024).
2. Ferdman, B. *et al.* VIPR: vectorial implementation of phase retrieval for fast and accurate microscopic pixel-wise pupil estimation. *Optics Express* **28**, 10179 (2020).
3. Nehme, E. *et al.* DeepSTORM3D: dense 3D localization microscopy and PSF design by deep learning. *Nature Methods* **17**, 734–740 (2020).
4. Mikhaylova, M. *et al.* Resolving bundled microtubules using anti-tubulin nanobodies. *Nat Commun* **6**, 7933 (2015).
5. Nieuwenhuizen, R. P. J. *et al.* Measuring image resolution in optical nanoscopy. *Nat Methods* **10**, 557–562 (2013).
6. Hoshen, Y., Li, K. & Malik, J. Non-adversarial image synthesis with generative latent nearest neighbors. in *Proceedings of the IEEE/CVF conference on computer vision and pattern recognition* 5811–5819 (2019).
7. Daniel, T. & Tamar, A. DDLP: Unsupervised Object-centric Video Prediction with Deep Dynamic Latent Particles. *Transactions on Machine Learning Research* (2024).
